# Genomic dissection of 43 serum urate-associated loci provides multiple insights into molecular mechanisms of urate control

**DOI:** 10.1101/743864

**Authors:** James Boocock, Megan Leask, Yukinori Okada, Asian Genetic Epidemiology Network (AGEN) Consortium, Hirotaka Matsuo, Yusuke Kawamura, Yongyong Shi, Changgui Li, David B Mount, Asim K Mandal, Weiqing Wang, Murray Cadzow, Anna L Gosling, Tanya J Major, Julia A Horsfield, Hyon K Choi, Tayaza Fadason, Justin O’Sullivan, Eli A Stahl, Tony R Merriman

**Author notes:** These authors contributed equally to the work (James Boocock, Megan Leask). These authors contributed equally to the work (Eli A Stahl, Tony R Merriman).

## Abstract

Serum urate is the end-product of purine metabolism. Elevated serum urate is causal of gout and a predictor of renal disease, cardiovascular disease and other metabolic conditions. Genome-wide association studies (GWAS) have reported dozens of loci associated with serum urate control, however there has been little progress in understanding the molecular basis of the associated loci. Here we employed trans-ancestral meta-analysis using data from European and East Asian populations to identify ten new loci for serum urate levels. Genome-wide colocalization with *cis*-expression quantitative trait loci (eQTL) identified a further five new loci. By *cis-* and *trans*-eQTL colocalization analysis we identified 24 and 20 genes respectively where the causal eQTL variant has a high likelihood that it is shared with the serum urate-associated locus. One new locus identified was *SLC22A9* that encodes organic anion transporter 7 (OAT7). We demonstrate that OAT7 is a very weak urate-butyrate exchanger. Newly implicated genes identified in the eQTL analysis include those encoding proteins that make up the dystrophin complex, a scaffold for signaling proteins and transporters at the cell membrane; *MLXIP* that, with the previously identified *MLXIPL*, is a transcription factor that may regulate serum urate via the pentose-phosphate pathway; and *MRPS7* and *IDH2* that encode proteins necessary for mitochondrial function. Trans-ancestral functional fine-mapping identified six loci (*RREB1, INHBC, HLF, UBE2Q2, SFMBT1, HNF4G*) with colocalized eQTL that contained putative causal SNPs (posterior probability of causality > 0.8). This systematic analysis of serum urate GWAS loci has identified candidate causal genes at 19 loci and a network of previously unidentified genes likely involved in control of serum urate levels, further illuminating the molecular mechanisms of urate control.

**Author Summary:** High serum urate is a prerequisite for gout and a risk factor for metabolic disease. Previous GWAS have identified numerous loci that are associated with serum urate control, however, only a small handful of these loci have known molecular consequences. The majority of loci are within the non-coding regions of the genome and therefore it is difficult to ascertain how these variants might influence serum urate levels without tangible links to gene expression and / or protein function. We have applied a novel bioinformatic pipeline where we combined population-specific GWAS data with gene expression and genome connectivity information to identify putative causal genes for serum urate associated loci. Overall, we identified 15 novel serum urate loci and show that these loci along with previously identified loci are linked to the expression of 44 genes. We show that some of the variants within these loci have strong predicted regulatory function which can be further tested in functional analyses. This study expands on previous GWAS by identifying further loci implicated in serum urate control and new causal mechanisms supported by gene expression changes.

## Introduction

Elevated serum urate (hyperuricemia) is causal of gout, an inflammatory arthritis increasing in prevalence world-wide [1, 2]. Monosodium urate crystals which form in hyperuricemic individuals can activate the NLRP3-inflammasome of resident macrophages to mediate an IL-1β-stimulated gout flare [3]. Long established genome-wide association studies (GWAS) [4, 5] have reported 28 loci associated with serum urate levels in European and East Asian sample sets with a more recent study reporting an additional 8 loci [6]. The loci of strongest effect are dominated by renal and gut transporters of urate, with two loci (*SLC2A9* and *ABCG2*) together explaining up to 5% of variance in serum urate levels in Europeans [4]. Most of these 36 loci also associate with gout in multiple ancestral groups [4, 7–9]. There has, however, been little progress on understanding the molecular basis of the association for the various loci. Probable causal genes have been identified at only about one fifth of the 36 loci [10–12], with strong evidence for causality for variants identified at *ABCG2* (*rs2231142*; Q141K) and *PDZK1* (*rs1967017*) [11, 13–17].

There have also been a number of recent improvements to the resources and analytical techniques that can be applied to the summary statistics of GWAS. Differences in underlying linkage disequilibrium (LD) structure between ancestral groups can be leveraged to amplify signal of association at shared causal variants [18, 19]. The epigenomics roadmap and ENCODE projects have generated a large resource of cell- and organ-specific regulatory regions [20, 21]. This information can be used to discover the cell-type specific regulatory regions that are known to be overrepresented in the heritability of a typical complex trait. Variants in regulatory regions identified by the epigenomics roadmap and ENCODE can be further analysed with functional annotation fine-mapping tools to identify candidate causal variants. Once credible sets of causal variants have been identified, expression quantitative trait loci (eQTL) sample sets (e.g. GTEx [22]) can be used to translate from causal variants to affected genes, thus informing the design of functional experiments for insights into molecular pathogenic pathways. Since sample sizes for eQTL studies are relatively modest (<1000), colocalization analyses of GWAS and eQTL data have remained primarily focused on *cis*-eQTL. However, recent methods that integrate high resolution genomic interaction data with eQTL data can reduce the number of *trans*-eQTL investigated substantially [23, 24], although a limitation of this filtering approach is that it excludes *trans*-eQTL not mediated by genomic interactions [25]. Despite this limitation integrating genomics interaction-filtered *trans*-eQTL signals with GWAS allows expansion of our view of how GWAS associations underpin gene expression [24].

In this study, we integrated these analytical approaches with the summary statistics of two serum urate GWAS from European and East Asian individuals [4, 5]. By meta-analysis and colocalization analysis of serum urate and eQTL signals we identified 15 new serum urate loci, identified 44 candidate causal genes connected to 25 loci, revealed the cell types that are enriched in serum urate heritability, and used this functional information to identify credible sets of causal variants using trans-ancestral fine-mapping.

## Results

### Trans-ancestral meta-analysis identifies 10 new loci associated with serum urate levels

The analysis approach for this study is summarised in Figure 1. Z-scores were imputed into the European [4] and East Asian [5] summary statistics using reference haplotypes from the Phase 3 1000 Genomes release and combined by meta-analysis (Figures 2 and S1). Study-specific results revealed three new loci at Chromosome 11 in the East Asian sample set (Chr11, 63.2-67.2Mb, *SLC22A9, PLA2G16, AIP*) in addition to those reported as genome-wide significant in the original GWAS [5] (Table 1; Figures 2, S2). All loci reported in the original GWAS reports [4, 5] were also detected in the trans-ancestral meta-analysis. However, the separate signals at *SLC22A11* and *SLC22A12* (Chr11, 64.4Mb) reported by Köttgen *et al*. [4] are reported as one signal in the trans-ancestral meta-analysis and an additional signal was detected at Chr11 65.4Mb (*RELA*). The trans-ancestral meta-analysis identified seven new loci (Chr4, 81.2Mb, *FGF5*; Chr5, 40.0Mb, *LINC00603*; Chr6, 32.7Mb, *HLA-DQB1*; Chr9, 33.2Mb, *B4GALT1*; Chr10, 60.3Mb, *BICC1*; Chr11, 63.9Mb, *FLRT1*; Chr11, 119.2Mb, *USP2*) (Figures 2 and S3). Of the ten new loci identified (seven from the trans-ancestral meta-analysis and three in the East Asian-specific analysis), five mapped within an extended Chr11 locus (63.2-67.2Mb) that encompassed the previously identified *SLC22A11, SLC22A12* and *OVOL1 / RELA* loci [4, 5]. In the East Asian GWAS, the peak marker falls outside the *RELA* locus (Figure S4). On closer inspection of the association signal from the region within and surrounding the *RELA* locus it is clear that the causal variants in the East Asian population are not the same as in the European population (Figure S4). On Chr6, given the association of the *HLA-DQB1* locus with T-cell-mediated autoimmunity [26] we also investigated if the lead *HLA-DQB1* SNP (*rs2858330*) was associated with other phenotypes using GWAS Central (www.gwascentral.org). There were no reported associations at *P* < 0.001, indicating that the *HLA-DQB1* signal in the serum urate GWAS is distinct from the association of this region with autoimmunity. The 35 loci found in Europeans explain 6.9% of variance in age and sex-adjusted serum urate levels. In summary, a total of 38 loci associated with serum urate concentration at a genome-wide level of significance were identified by this analysis.

**Figure 1.**
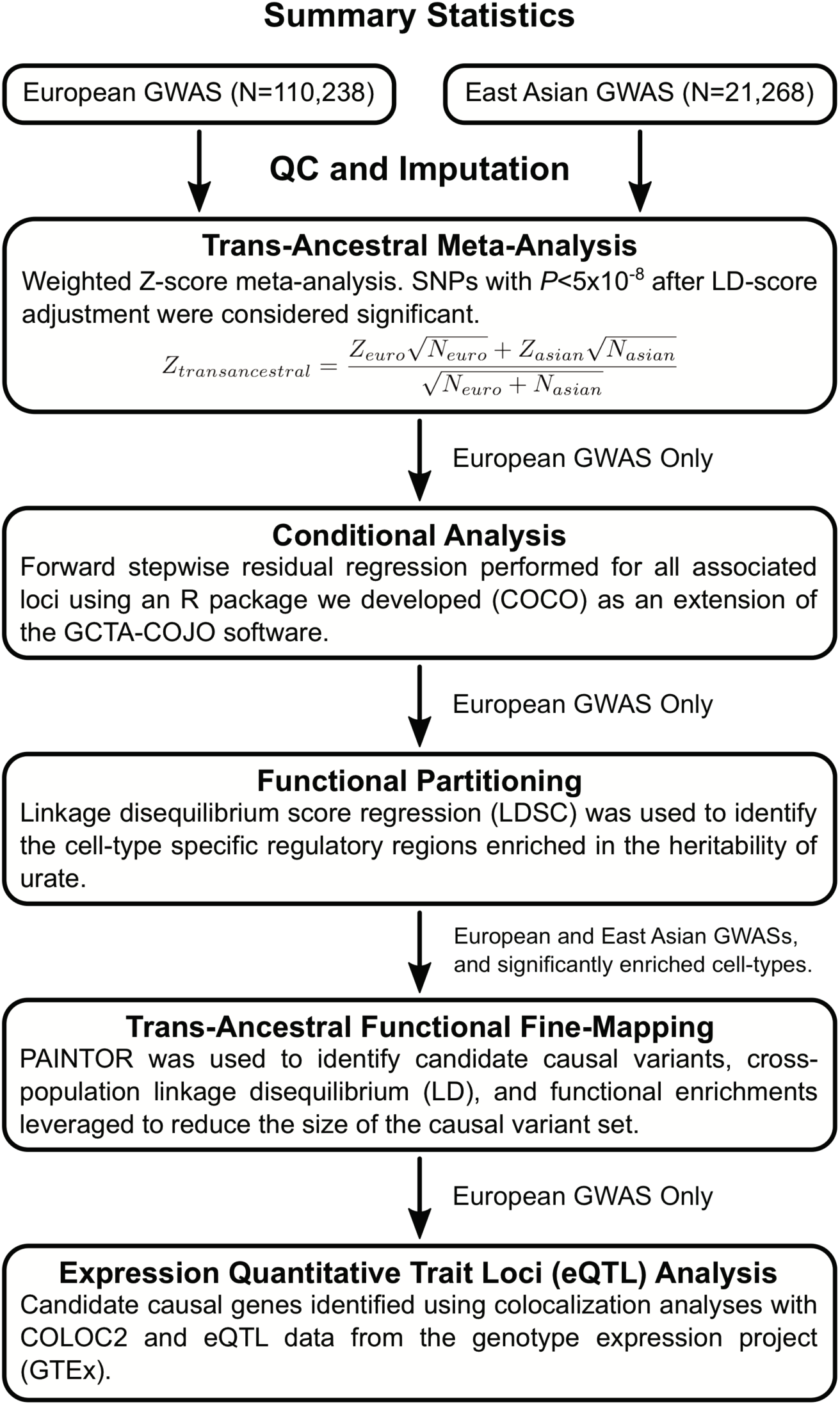
Analysis flowchart.

**Figure 2.**
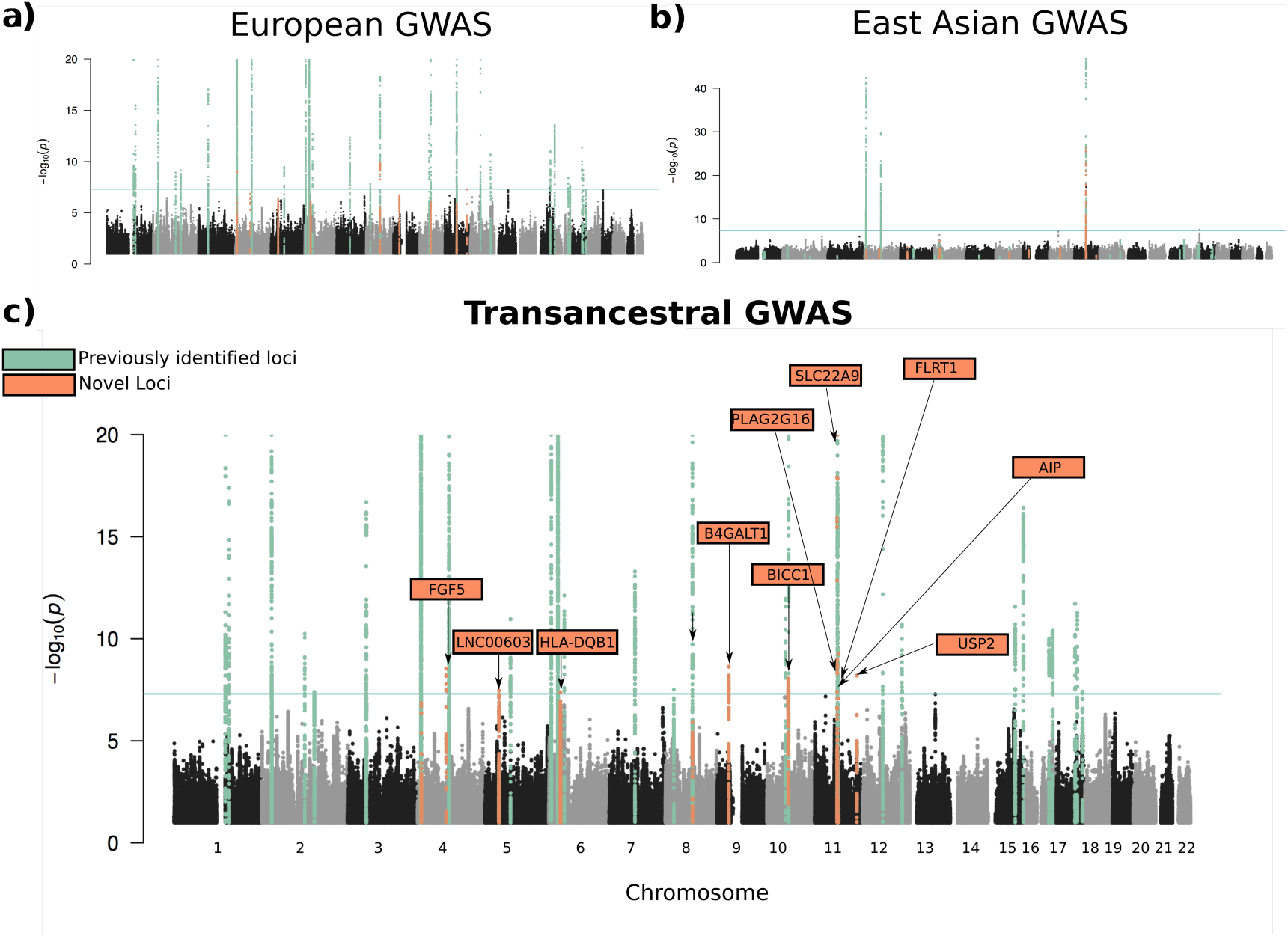
Manhattan plots showing -log_10_(*P*) for all SNPs of the European, East Asian, and trans-ancestral GWAS ordered by chromosomal position. (A) Manhattan plot of the European GWAS. (B) Manhattan plot of the East Asian GWAS. (C) Manhattan plot of the trans-ancestral GWAS. SNPs within previously identified serum urate loci are colored light green. SNPs located within novel serum urate loci are colored orange. For the ten new genome-wide significant loci identified by trans-ancestral meta-analysis, the closest gene to the lead SNP is indicated.

**Table 1.**
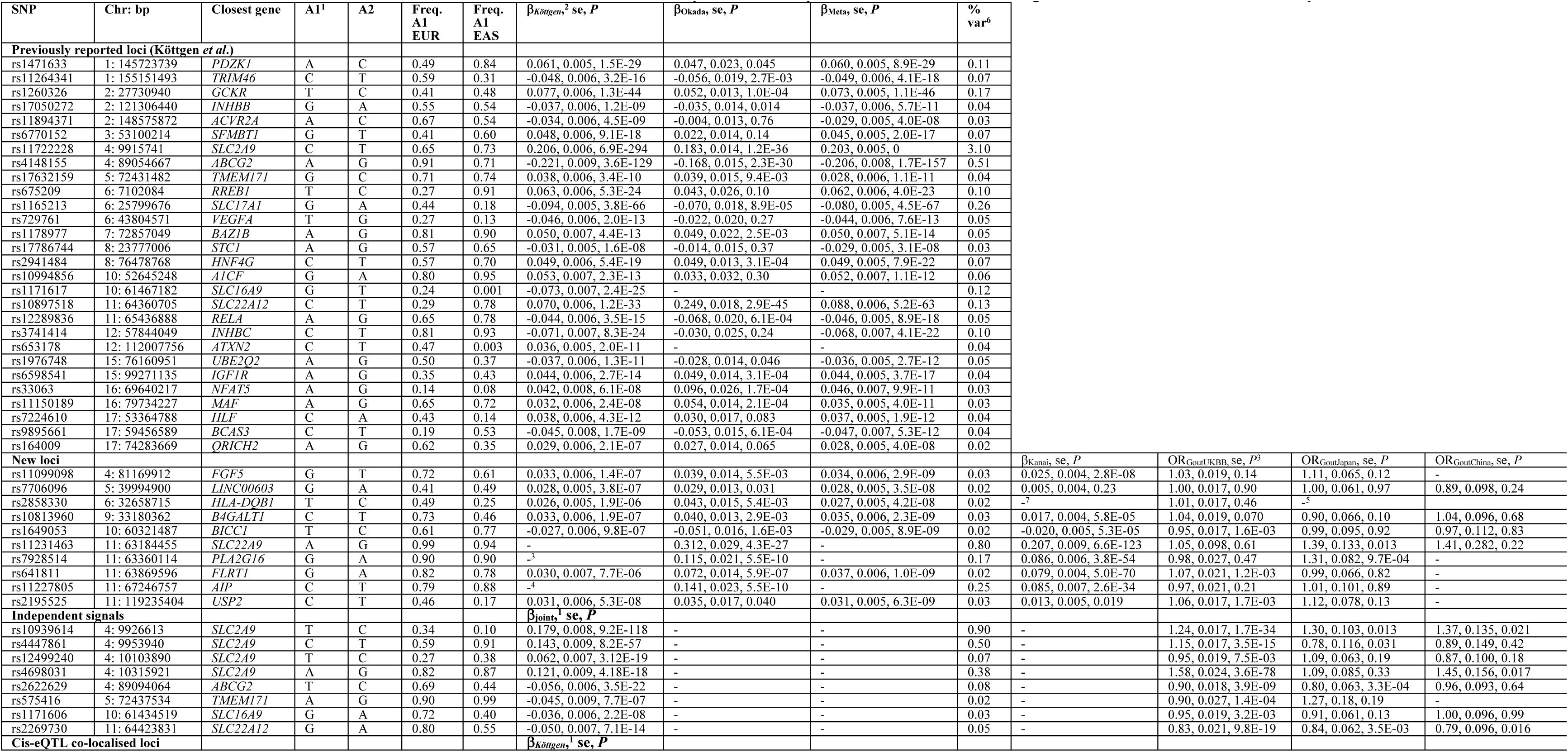
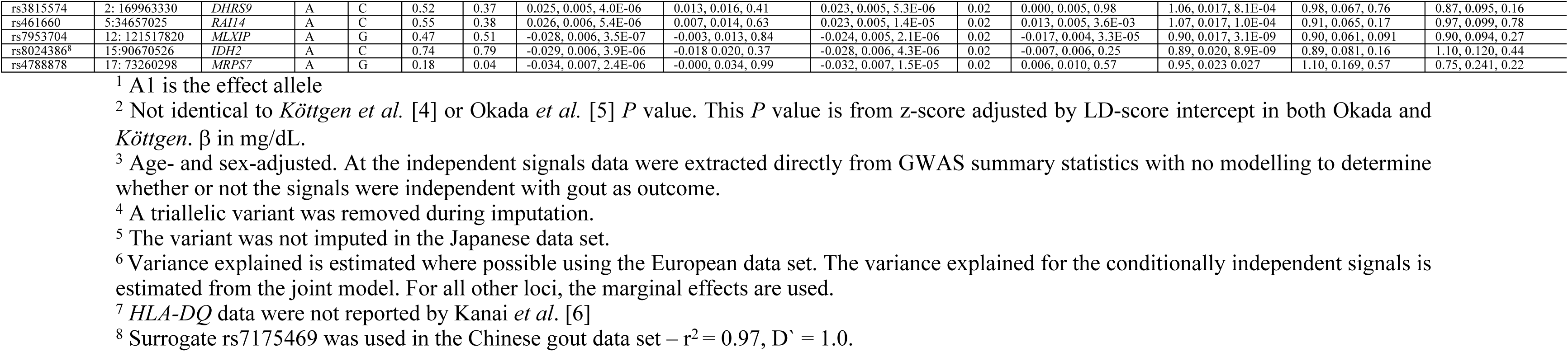
SNPs associated with serum urate concentrations by meta-analysis individuals of European and East Asian ancestry.

### Conditional analysis identifies 8 additional variants associated with serum urate

Using the European summary statistics, a conditional and joint analysis was performed with the objective of identifying independent genetic effects. Conditional and joint analysis identified an additional four genome-wide significant associations at *SLC2A9*, and one at each of *ABCG2, TMEM171, SLC16A9* and *SLC22A11/A12* (Table 1). The conditional analysis was limited to four independent associations at each locus, therefore it remains possible that there are additional unidentified associations at these loci. In a joint model these four loci explained an additional 0.54% of the variance of age- and sex-adjusted serum urate levels (Table 1).

### Population-specific associations with serum urate levels

LocusZoom plots from each population were visually compared to the trans-ancestral meta-analysis to identify population-specific and shared patterns of association. For 16 loci (*ABCG2, B4GALT1, BCAS3, FGF5, BICC1, HFN4G, IGF1R, INHBB, NFAT5, PDZK1, QRICH2, SLC16A9, SLC17A1, TMEM171, TRIM46, UBE2Q2*) the pattern of association was consistent between the East Asian and European GWAS suggesting strong similarity between the underlying haplotypic structure and casual variant(s) (Figure S5). The *MAF* locus contains two association signals in the East Asian population, one that is shared with the European population and one that is specific to the East Asian population (Figure S6) [12]. The lead SNP for the East Asian population is *rs889472*; this variant is common in both European (C-allele = 0.38) and East Asian (C-allele = 0.60) individuals from the 1000 Genomes Project yet there is no serum urate association signal in the European population. Two other East Asian-specific signals were identified on chromosome 11 near the *SLC22A9* and *PLA2G16* genes in addition to the previously mentioned East Asian-specific signal at the *RELA* locus. These loci, in combination with the conditionally independent and trans-ancestral associations, mean that there are seven independent associations on chromosome 11 between 63.1Mb and 67.3Mb.

### Cis-eQTL colocalization analysis identifies 24 candidate causal genes at 19 serum urate loci

To connect the serum urate associations with the genes they influence, we utilised publicly available expression data provided by the GTEx consortium and performed colocalization with COLOC [27] (Figure S7 and Table 2). This method attempts to identify whether the causal variant is the same in both the eQTL and GWAS signal indicating a putative causal mechanism, whereby the variant alters gene expression (transcript levels) and expression influences the trait – in this case serum urate levels. This approach provides further support for the loci identified by the trans-ancestral meta-analysis by linking the serum urate signals into the biological process of gene regulation. For 19 of the serum urate GWAS loci strong evidence for colocalization (PPC > 0.8) was seen with 24 *cis*-eQTL (Table 2; Figure S7). The 19 loci included five loci identified by inclusion of sub-genome-wide significant GWAS loci in the analysis (*DHRS9, RAI14, MLXIP, IDH2, MRPS7*). For 11 of the previously identified Köttgen *et al*. [4] GWAS loci there are colocalized *cis*-eQTL (*PDZK1, TRIM46, INHBB, SFMBT1, BAZ1B, SLC16A9, INHBC, UBE2Q2, IGF1R, MAF, QRICH2*). Of the ten new loci discovered as genome-wide significant in the trans-ancestral meta-analysis colocalized eQTL were identified at three loci (*HLA-DQB1, B4GALT1, RELA*).

**Table 2.** Serum urate associated loci with colocalized GTEx eQTL.

| Locus | Lead GWAS variant | Colocalized eQTL gene | PPC<sup>1</sup> | Tissue(s) | Direction (allele, $\beta_{SU}$ mg/dL, $\beta$ Expression, $P$ Expression) |
| --- | --- | --- | --- | --- | --- |
| Cis-eQTL (genome-wide significant by trans-ancestral meta-analysis) |  |  |  |  |  |
| *PDZK1* | rs1471633 | *PDZK1* | 0.98 | colon - transverse, small intestine | A, 0.061, 0.58, 1.3E-11 (colon - transverse) |
| *TRIM46* | rs11264341 | *MUC1* | 0.93 | adipose subcutaneous, artery aorta, esophagus mucosa, esophagus muscularis, testis, whole blood | T, -0.048, 0.34, 7.7E-16 (esophagus – mucosa) |
|  |  | *GBAP1* | 0.98 | skin | T, -0.048, -0.35, 2.7E-08 |
|  |  | *FAM189B* | 0.92 | heart atrial appendage | T, -0.048, -0.23, 1.1E-05 |
| *INHBB* | rs17050272 | *INHBB* | 0.80 | lung | G, -0.037, 0.23, 2.2E-05 |
| *SFMBT1* | rs6770152 | *TMEM110* | 0.86 | adipose subcutaneous, skin | T, -0.048, 0.24, 1.1E-05 (adipose subcutaneous) |
|  |  | *SFMBT1* | 0.97 | colon transverse | T, -0.048, 0.45, 1.9E-10 |
| *HLA-DQB1* | rs2858330 | *HLA-DQA2* | 0.82 | prostate | T, 0.026, -0.79, 2.8E-12 |
| *BAZ1B* | rs1178977 | *MLXIPL* | 0.88 | adipose visceral, transformed fibroblasts | T, 0.050, -0.48, 1.2E-08 (transformed fibroblasts) |
| *B4GALT1* | rs10813960 | *B4GALT1* | 0.81 | EBV transformed lymphocytes, esophagus mucosa | T, -0.033, -0.47, 1.9E-04 (EBV transformed lymphocytes) |
| *SLC16A9* | rs1171617 | *SLC16A9* | 0.84 | artery aorta, thyroid | T, 0.073, -0.28, 2.2E-06 (thyroid) |
| *RELA* | rs12289836 | *OVOL1-AS1* | 0.88 | thyroid, caudate basal ganglia, cortex | A, -0.043, 0.55, 3.7E-06 (basal ganglia)<sup>2</sup> |
| *INHBC* | rs3741414 | *R3HDM2* | 0.84 | transformed fibroblasts | T, -0.071, 0.21, 2.2E-05 |
| *UBE2Q2* | rs1976748 | *UBE2Q2* | 0.91 | dorsolateral prefrontal cortex | A, -0.037, -0.10, 8.5E-30 |
| *IGF1R* | rs6598541 | *IGF1R* | 0.83 | heart left ventricle | A, 0.044, -0.33, 1.7E-07 |
| *MAF* | rs11150189 | *MAFTRR* | 0.84 | colon sigmoid, pancreas | A, 0.032, 0.52, 1.2E-05 (colon sigmoid) |
| *QRICH2* | rs164009 | *UBALD2* | 0.90 | esophagus muscularis, caudate basal ganglia, whole blood | A, 0.029, 0.23, 7.7E-07 (esophagus muscularis) |
|  |  | *PRPSAP1* | 0.85 | anterior cingulate cortex | A, 0.029, 0.62, 1.7E-06 |
| **Cis-eQTL (sub genome-wide significant by trans-ancestral meta-analysis)** |  |  |  |  |  |
| *DHRS9* | rs3815574 | *DHRS9* | 0.83 | whole blood | A, 0.024, -0.30, 1.7E-26 |
| *RAI14* | rs461660 | *RAI14* | 0.83 | thyroid | A, 0.026, 0.24, 3.5E-07 |
| *MLXIP* | rs7953704 | *MLXIP* | 0.83 | small intestine | A, -0.028, 0.48, 9.6E-07 |
| *IDH2* | rs8024386 | *IDH2* | 0.87 | atrial appendage | A, -0.029, 0.35, 1.1E-06 |
| *MRPS7* | rs4788878 | *GGA3*<br>*MRPS7* | 0.83<br>0.87 | thyroid<br>dorsolateral prefrontal cortex, colon transverse, transformed fibroblasts, pancreas | A, -0.034, 0.23, 1.9E-08<br>A, -0.034, 0.07, 2.5E-12 (dorsolateral prefrontal cortex) |
| **Trans-eQTL** |  |  |  |  |  |
| *NFAT5* | rs33063 | *AIF1L* | 0.92 | brain substantia nigra | A, 0.042, 0.39, 1.9E-05 |
|  |  | *CACNA2D3* | 0.99 | basal ganglia | A, 0.042, 0.23, 4.2E-06 |
|  |  | *STIM1* | 0.99 | basal ganglia | A, 0.042, 0.26, 5.6E-07 |
| *SLC16A9* | rs1171617 | *ANKS1B* | 0.98 | testis | T, 0.073, 0.17, 1.8E-05 |
|  |  | *DSCAM* | 0.98 | brain hypothalamus | T, 0.073, 0.51, 1.9E-05 |
| *BAZ1B* | rs1178977 | *RNF24* | 0.98 | brain cortex | A, 0.050, -0.43, 1.3E-05 |
| *QRICH2* | rs164009 | *PPP3R1* | 0.98 | heart left ventricle | A, 0.029, -0.18, 7.0E-06 |
| *INHBB* | rs17050272 | *CHAC2* | 0.97 | basal ganglia | A, 0.037, -0.36, 1.3E-05 |
|  |  | *ZNF804A* | 0.99 | brain anterior cingulate cortex | A, 0.037, 0.23, 1.1E-05 |
| *UBE2Q2* | rs1976748 | *COL11A1* | 0.95 | colon transverse | A, -0.037, 0.23, 6.0E-06 |
| *HNF4G* | rs2941484 | *CSMD2* | 0.99 | brain cerebellum | T, 0.049, -0.40, 5.2E-06 |
| *INHBC* | rs3741414 | *SPIN1* | 0.98 | brain frontal cortex | T, -0.071, -0.22, 1.7E-05 |
| *DHRS9* | rs3815574 | *JHDM1D* | 0.94 | brain hippocampus | A, 0.024, 0.32, 1.3E-05 |
| *IDH2* | rs8024386 | *MAPK6* | 0.86 | brain amygdala | A, -0.029, -0.32, 1.2E-04 |
|  |  | *ZBTB20* | 0.89 | testis | A, -0.029, 0.10, 1.8E-05 |
| *RREB1* | rs675209 | *UTRN* | 0.99 | brain putamen basal ganglia | T, 0.063, 0.35, 2.1E-05 |
| *HLF* | rs7224610 | *DMD* | 0.94 | brain nucleus accumbens basal ganglia | A, -0.038, -0.36, 1.9E-05 |
| *VEGFA* | rs729761 | *CLPS* | 1.00 | brain cerebellar hemisphere | T, -0.046, -0.59, 3.8E-06 |
| *MLXIP* | rs7953704 | *NDUFA12* | 0.95 | brain putamen basal ganglia | A, -0.028, -0.32, 2.0E-05 |
| *BCAS3* | rs9895661 | *TMEM117* | 0.99 | prostate | T, 0.045, 0.48, 3.9E-06 |

<sup>1</sup> Posterior probability of colocalization (PPC)
<sup>2</sup> Data from proxy variant rs642803.

### CoDeS3D analysis integration with GTEx and colocalization for identification of trans-eQTL

To identify candidate causal genes that represent *trans*-eQTL, we pre-screened for SNP-gene physical connectivity using the CoDeS3D algorithm and then tested for colocalization with serum urate GWAS signals (Table 2). This identified 20 *trans*-eQTL signals that co-localized (PPC > 0.8) with 15 GWAS loci (Figure S7). Of the 20 genes with colocalized *trans-*eQTL we identified, only two had evidence within the gene (*P* < 5 x 10 ^−04^) for a signal of association with serum urate by GWAS (Figure S8) – *UTRN* in the Köttgen *et al*. dataset (lead variant *rs4896735*, *P* = 2 x 10^−04^) and *DMD* (*rs1718043*; *P* = 9 x 10^−05^) in the Kanai *et al*. [6] dataset. The *DMD* and *UTRN* genes encode components of the dystrophin complex. Notably, *MAPK6* (also known as *ERK3*) and a *trans*-eQTL identified at the *IDH2* locus has a signal of association with serum urate levels in response to allopurinol in gout by GWAS (*rs62015197*, *P* = 8 x 10^−07^) [28].

Nine serum urate loci (*SLC16A9, BAZ1B, QRICH2, UBE2Q2, INHBB, INHBC, DHRS9, MLXIP, IDH2*) exhibited both *cis-* and *trans*-eQTL of which the latter three had been identified by the genome-wide colocalization analysis. At *SLC16A9*, the signal is different between the *cis-* and *trans*-eQTL (Figure S7), with all of the GWAS signal present in the *cis*-eQTL whereas only the signal associated with the lead GWAS SNP was evident in the *trans*-eQTL. Also, at *SLC16A9* there was a second *cis*-eQTL over *CCDC6* that was weakly associated with serum urate levels. Differential *cis-* and *trans*-eQTL signals are reminiscent of the situation at the serum urate-associated *cis-* and *trans*-eQTL signals at the *MAFTRR* locus [12].

### Replication in Kanai et al

While this work was being finalized a serum urate GWAS comprising 109,029 Japanese individuals (of whom 18,519 over-lapped with the Okada *et al*. [5] study) was published [6] allowing an opportunity to replicate our findings. Seven of the 15 new loci we identified replicated (*P* < 0.003) in the Kanai *et al*. [6] study (Table 1). The replicated loci included two (*FGF5* and *BICC1*) of the total 27 genome-wide significant signals reported by Kanai *et al*. – of the remaining 25 loci identified by Kanai *et al*. [6] 17 had previously been reported by others [4, 29, 30] and eight new (the gene containing the lead SNP or the flanking genes at each locus: *RNF115* (*rs12123298*), *USP23* (*rs7570707*), *UNCX* (*rs4724828*), *TP53INP1* (*rs7835379*), *EMX2/RAB11F1P2* (*rs1886603*), *SBF2* (*rs2220970*), *MPPED2/DCDC5* (*rs963837*), *GNAS* (*rs6026578*)).

### Testing for association with gout

To replicate the urate signals we tested the independent signals at eight existing loci, ten new loci with genome-wide significance in the trans-ancestral meta-analysis and five loci discovered by colocalization with eQTL (Table 1) for association with gout in European (UK Biobank) [31], Chinese [32] and Japanese [30] sample sets. The *BICC1, FLRT1* and *USP2* loci replicated (*P ≤* 1.6 x 10^−03^) in the European dataset in a directionally-consistent fashion (i.e. the urate-increasing allele associated with an increased risk of gout). The *SLC22A9* and *PLA2G16* loci replicated (*P ≤* 0.013) in the Japanese dataset also in a directionally-consistent fashion. All eight additional variants identified in the European serum urate data set by conditional analysis (Table 1) were replicated (*P ≤* 3.3 x 10^−03^), in the European gout data set. For the five loci identified by colocalization with eQTL, all replicated (*P ≤* 0.027) in the European gout data set, with *IDH2* and *MLXIPL* at a genome-wide level of significance (*P* < 5.0 x 10^−08^). All had an OR for gout consistent with the direction of effect on serum urate levels. None of these five loci were associated with gout in the Chinese or Japanese sample sets.

### Functional partitioning of the heritability of serum urate levels

To understand the functional categories that contribute most to the heritability of serum urate level, we used LD score regression to functionally partition the SNP heritability of the European serum urate GWAS (Figures 3, S9; Table S1). Functional partitioning of serum urate SNP heritability according to cell type revealed significant enrichments in the kidney (*P* = 3.2 x 10^−08^), the gastrointestinal tract (*P* = 5.2 x 10^−08^), and the liver (*P* = 3.4 x 10^−03^). A refined analysis of 218 functional annotations, which contribute to the larger cell type groups, revealed 11 significant annotations: four histone marks in the kidney H3K27ac (*P* = 1.2 x 10^−07^), H3K9Ac (*P* = 1.5 x 10^−06^), H3K4me3 (*P* = 9.6 x 10^−0.6^), and H3K4me1 (*P* = 2.5 x 10^−05^) and two histone marks in the gastrointestinal tract - H3K27ac (*P* = 5.6 x 10^−06^), and H3K4me1 (*P* = 4.8 x 10^−05^). These histone marks are characteristic of transcriptional activation and consistent with active expression of nearby genes.

**Figure 3.**
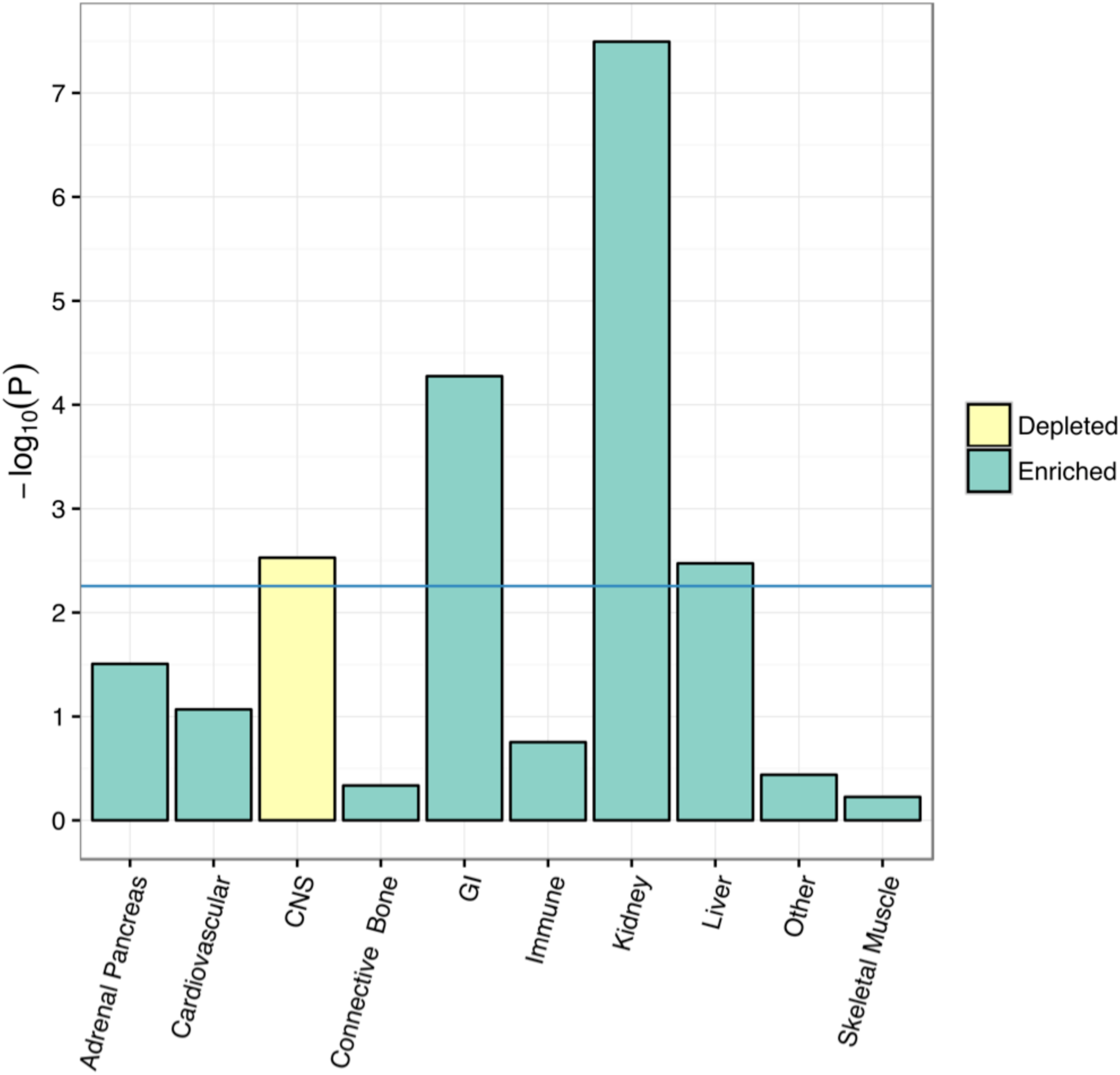
Tissue-focused functional heritability enrichments. Tissue-focused functional heritability enrichments for serum urate levels. The color of each bar indicates whether heritability was depleted or enriched within a particular cell-type group. The –log10 P-value for the enrichment in each cell-type group is on the Y-axis. These enrichments were generated using LD-Score functional partitioning of the European GWAS summary statistics.

### Trans-ancestral functional fine-mapping identifies putative causal variants

We sought to leverage both the functional enrichments and linkage disequilibrium differences between the populations to identify candidate causal variants at each locus associated with serum urate levels. To this end, we performed trans-ancestral fine-mapping with PAINTOR using the kidney, gastrointestinal tract, and liver cell type group annotations as functional priors. When analysing only the European GWAS the 90% causal credible sets had on average 129 SNPs. With the addition of the East Asian GWAS data the set size reduced to an average of 56 SNPs, and functional annotations reduced the average credible set size to 41. Of the 36 loci used in this analysis (the 28 reported by Köttgen *et al.* [4] and the ten new genome-wide significant loci reported here, excluding *RELA* and *HLA-DQB1*), 14 loci had seven or fewer causal variants in their 80% causal credible set (Tables 3, S3). The combination of both the functional annotations and East Asian GWAS data significantly improves our ability to identify the causal variants for loci associated with serum urate levels.

**Table 3.** Putative credible causal SNP set identified with PAINTOR.

| Locus | Chr:pos | SNP ID | Posterior prob | European Z-score<sup>2</sup> | East Asian Z-score | Motifs changed (haploreg) |
| --- | --- | --- | --- | --- | --- | --- |
| SLC2A9 | 4:9915741 | rs11722228 | 1.000 | -36.73 | -12.67 | None |
| SLC2A9 | 4:9946095 | rs4697701 | 1.000 | 59.52 | 9.52 | E2a, Mxi1 |
| SLC2A9 | 4:9954660 | rs11723382 | 1.000 | 38.02 | 7.24 | Hmx, Nkx2 |
| SLC2A9 | 4:9981997 | rs13145758 | 1.000 | -57.11 | -2.66 | Egr1, GCNF, HNF4 |
| SLC2A9 | 4:9982330 | rs13125646 | 1.000 | -49.28 | -2.66 | None |
| ABCG2 | 4:88917735 | rs17013705 | 1.000 | 4.44 | 5.10 | Irf, TATA, TCF12 |
| ABCG2 | 4:88944511 | rs2725227 | 1.000 | -7.22 | -2.52 | Gfi1, TCF11, MafG |
| ABCG2 | 4:88960528 | rs2725217 | 1.000 | -17.56 | -9.73 | CDP7, Dbx1, HNF1, Mef2, Pouf1, TATA |
| ABCG2 | 4:88973427 | rs2725210 | 1.000 | -15.27 | -4.69 | Fox, FoxA, FoxC1, FoxJ2, FoxF2, FoxK1, PLZF |
| ABCG2 | 4:88999222 | rs2728126 | 1.000 | -15.65 | -8.72 | EWRS1, FLI1 |
| ABCG2 | 4:89052323 | rs2231142 | 1.000 | -24.26 | -11.43 | GR, Irf |
| ABCG2 | 4:89098731 | rs9631715 | 1.000 | 15.34 | 7.24 | GATA, SREBP |
| SLC17A1 | 6:25785295 | rs6909187 | 1.000 | 15.65 | -0.77 | FoxC1, HDAC2, HMG-IY, Pou2F2 |
| SLC17A1 | 6:25786993 | rs3799344 | 1.000 | 15.36 | 2.10 | Eomes, Pax6, TBX5 |
| UBE2Q2 | 15:76194286 | rs335685 | 1.000 | 5.25 | 0.13 | CEBPB, DMRT1, FoxA, Nanog, Nkx6, Pou2F2, Pou3F4, STAT, TATA |
| SFMBT1 | 3:53092375 | rs9870898 | 0.996 | 5.15 | 0.63 | AhR, GR, HES1, HNF1, Pax4 |
| SFMBT1 | 3:53026384 | rs2564938 | 0.996 | -6.86 | -1.46 | None |
| SFMBT1 | 3:53026714 | rs2115779 | 0.996 | 5.38 | 1.63 | Hsf, Ptf1b |
| SLC22A12 | 11:64333296 | rs1783811 | 0.994 | 7.91 | 5.00 | Mef2, TAL1, ZID |
| UBE2Q2 | 15:76160951 | rs1976748 | 0.991 | -6.78 | -2.00 | Arid3a, Sox, TCF4 |
| AIP | 11:67246757 | rs11227805 | 0.991 | NA<sup>1</sup> | 6.22 | None |
| SLC22A12 | 11:64358241 | rs11602903 | 0.990 | 11.16 | 13.45 | BAF155 |
| SLC22A12 | 11:64387932 | rs2277311 | 0.990 | 10.79 | 14.29 | Hic1 |
| SLC22A12 | 11:64338228 | rs11231822 | 0.990 | 12.88 | -2.62 | CTCF, ERalphaA, Lmo2, Nanog |
| SLC22A12 | 11:64419217 | rs502571 | 0.990 | -8.41 | -10.70 | Mrg1, Hoxa9, TAL1 |
| SLC22A12 | 11:64474752 | rs2957564 | 0.990 | -8.45 | -10.94 | GR, LUN1 |
| SLC22A12 | 11:64622502 | rs2007521 | 0.990 | -1.50 | -6.90 | AP1, CTCF, Ets |
| SLC17A1 | 6:25798932 | rs1165215 | 0.986 | -16.73 | -3.92 | None |
| SLC17A1 | 6:26125342 | rs129128 | 0.986 | 5.24 | -0.09 | None |
| SLC22A9 | 11:63170736 | rs7925182 | 0.981 | -1.26 | -0.94 | Barx1 |
| USP2 | 11:119235404 | rs2195525 | 0.978 | 5.46 | 2.06 | AP1, Foxa, STAT |
| RREB1 | 6:7102084 | rs675209 | 0.964 | 10.13 | 1.63 | CCNT2, Ets, MZF1, NRSF, STAT, VDR, Zfp281, Zfp740 |
| HNF4G | 8:76401359 | rs13264750 | 0.964 | 0.44 | -2.78 | Pou3f2 |
| HLF | 17:53364788 | rs7224610 | 0.920 | 6.95 | 1.74 | None |
| GCKR | 2:27730940 | rs1260326 | 0.868 | 14.00 | 3.89 | NRSF |
| INHBC | 12:57807114 | rs540730 | 0.851 | -9.44 | 0.75 | None |
| SLC22A9 | 11:63859120 | rs11231454 | 0.810 | -0.31 | 10.01 | GATA |
| SLC22A9 | 11:63171309 | rs12281229 | 0.810 | -1.01 | -3.09 | Cdx, Dbx1, Fox, FoxA, fOxc1, FoxD3, FoxF1, FoxI1, FoxJ1, Foxj2, FoxK1, FoxL1, Foxo, FoxP1, HDAC2, HNF1, Hlx1, HoxD8, Mef2, NF-Y, Ncx, Pbx-1, Pbx3, TATA |

<sup>1</sup> At AIP
<sup>2</sup> Z-scores are reported because effect sizes are not available for imputed variants.

*SLC2A9* is a complex locus with a very strong effect on serum urate levels and multiple independent genetic effects [33, 34]. A subset of the lead urate SNPs at *SLC2A9* with PAINTOR posterior probabilities of 1.0 overlap putative regulatory elements (Tables 3, S4). One of these urate-associated variants at *SLC2A9*, *rs11723382*, is also among the maximally-associated *cis*-eQTL variants for *RP11-448G15.1* (transformed lymphocytes) (*RP11-448G15.1* is a lncRNA located within the second intron of *SLC2A9*) and disrupts two predicted motifs Hmx and Nkx2 (Figure S10 and Table 3) [35]. This eQTL was not identified in our COLOC analysis, however visual inspection of the *RP11-448G15.1* eQTL and *SLC2A9* GWAS signal indicates that the signals coincide and suggests *RP11-448G15.1* expression is likely important for serum urate control.

At *SLC22A12 / NRXN2*, four of the seven putative causal variants (Table 3) are in LD (R^2^ > 0.6) with the maximal *trans*-eQTL variant for *RNF169* identified by CoDeS3D. Visual inspection of the *RNF169 trans*-eQTL and the serum urate signal at the *SLC22A12 / NRXN2* locus indicates that these signals overlap (Figure S10). *rs2277311*, an intronic variant located within *NRXN2*, is the most likely candidate of these variants to have regulatory function. *rs2277311* has promoter, enhancer and DNase signatures and the urate-decreasing A-allele disrupts a predicted HiC1 motif (Tables 3, S4) [35].

Six loci (*RREB1, INHBC, HLF, UBE2Q2, SFMBT1, HNF4G*) with PAINTOR causal SNPs (PP > 0.8) also have colocalised eQTL (Tables 2 and 3). These loci represent good candidates for follow up analyses of regulatory function (e.g. [11, 12]). The lead urate variant at the *HLF* locus, *rs7224610* (PAINTOR posterior probability = 0.92) is intronic, has enhancer signatures, is bound by multiple transcription factors including POL2 (Table S4) and is amongst the maximally associated *trans*-eQTL variants for *DMD* (encodes *dystrophin*). *rs675209* at *RREB1* is the maximal *trans*-eQTL variant for *UTRN* (encodes *utrophin*), overlaps enhancer signatures in six tissues and alters 8 transcription factor binding motifs (Tables 3 and S4). The variants at *HNF4G*, *SFMBT1, UBE2Q2* and *INHBC* do not overlap putative regulatory elements (Table S4). Although *rs13264750 (HNF4G)*, *rs2115779 (SFBMT1)* and *rs9870898* (upstream of *SFMBT1*) are predicted to change 8 binding motifs including HNF1 (Table 3).

### SLC22A9

*SLC22A9* encodes organic anion transporter 7 (OAT7). OAT7, expressed only in the liver, is a relatively poorly characterized member of the OAT family [36] that includes urate secretory transporters OAT1-3 and the urate reuptake transporter OAT4 (encoded by *SLC22A11*) [37]. RT-PCR screening of human cell lines indicated expression in HepG2 cells (Figure 4). OAT7 exhibited modest uricosuric-sensitive urate uptake when expressed in *Xenopus* oocytes (Figure 4). Pre-injection of oocytes with butyrate, but not other anions (data not shown), led to a modest trans-activation of urate transport, consistent with urate-butyrate exchange.

**Figure 4.**
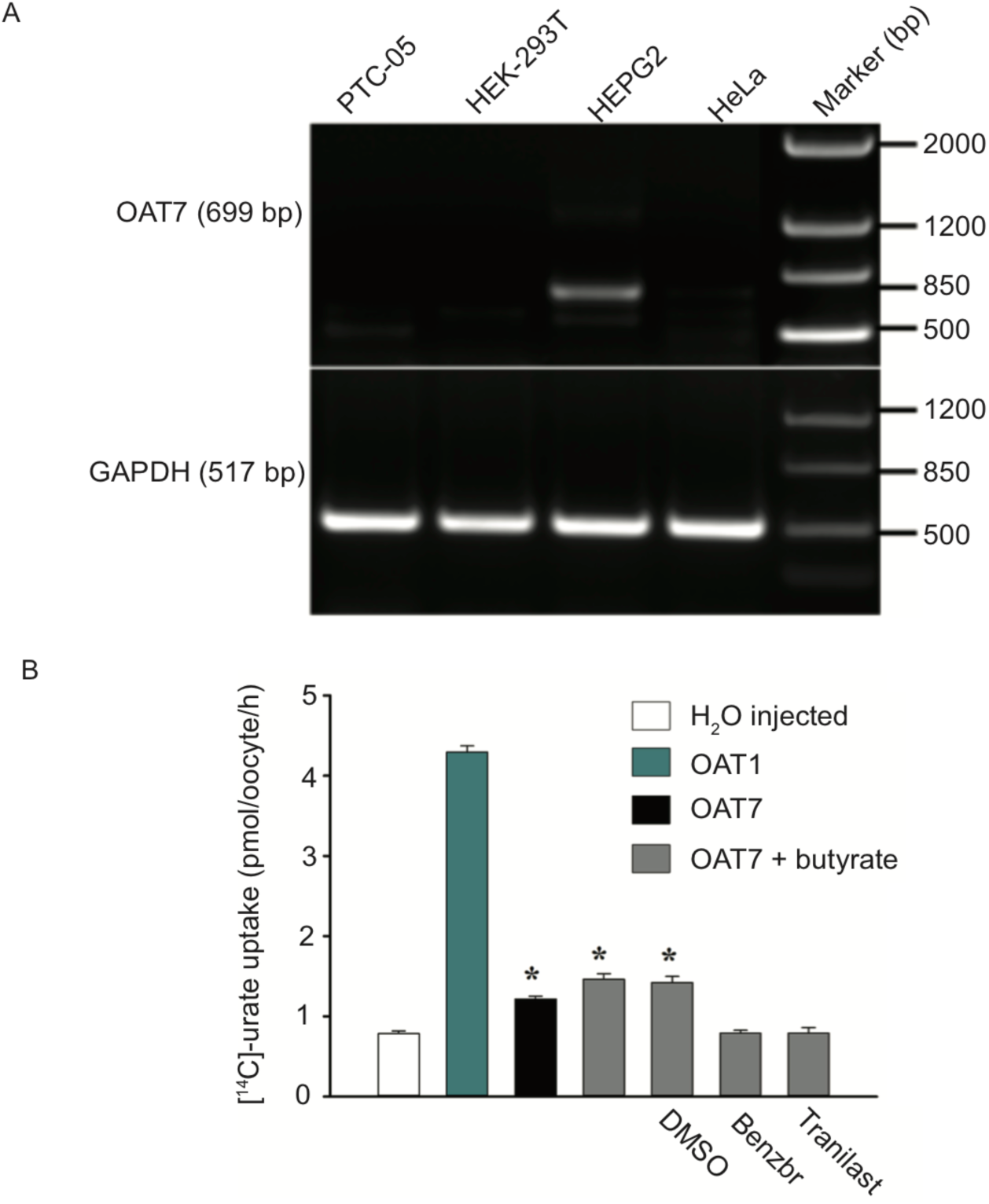
Expression analysis and functional expression of *SLC22A9* (OAT7). (A) RT-PCR of *SLC22A9*/OAT7 expression in the human PTC-05 proximal tubular cell line, HEK-293T cells, and HepG2 hepatic cells. All three cell lines are positive for GAPDH but *SLC22A9*/OAT7 is unique to HepG2. (B) OAT7 is a weak urate transporter. *Xenopus* oocytes were microinjected with water (control cells) or cRNA for OAT1 or OAT7. OAT7-expressing cells have a very modest urate transport activity that is increased by prior microinjection with butyrate, to “trans-activate” urate-butyrate exchange. This transport activity is inhibited by the uricosurics tranilast and benzbromarone, each at a concentration of 100 µM; DMSO, the diluent for tranilast and benzbromarone, has no effect on urate transport. * refers to *P*<0.001 compared to OAT7-expressing cells without butyrate pre-injection and water control cells. Data shown are from a single representative experiment.

## Discussion

### Identification of 15 new loci associated with serum urate

The 15 new loci identified here as associated with serum urate levels can be ranked according to the strength of genetic evidence according to two criteria; a genome-wide significant association with serum urate levels, replication in gout, and/or replication in the recently published Japanese serum urate GWAS [6]. In addition to the strength of the genetic evidence, seven of these loci were co-localized with at least one eQTL signal which identifies a putative causal gene, and provides further evidence of a genuine association with serum urate levels. Of the 15 novel loci, eight (*FGF5, BICC1*, *PLA2G16, B4GALT1, SLC22A9, AIP, FLRT1* and *USP2*) were genome-wide significant and replicated in gout (Table 1) or the Kanai *et al*. urate dataset [6]. The *HLA-DQB1* locus was genome-wide significant and a putative causal gene *HLA-DQA2* was identified. The *DHRS9, MLXIP, MRPS7, RAI14,* and *IDH2* loci were only of suggestive association in the trans-ancestral meta-analysis but the colocalization analysis provided strong evidence that they participate in a causal pathway. Of these five loci, we were able to replicate the association at *DHRS9*, *MLXIP, MRPS7, RAI14* and *IDH2* in gout, and for *MLXIP* we additionally replicated the association in the Kanai *et al*. [6] data. Overall, the evidence that these five loci, identified solely by colocalization of GWAS signal with an eQTL signal, have a true association with serum urate is strong and provide empirical support for our genome-wide co-localization approach using sub-genome wide significant GWAS signals. Overall, we identified 14 novel loci that we are confident are unlikely to represent false positive associations (*FGF5*, *B4GALT1, PLA2G16, SLC22A9, FLRT1, USP2, BICC1, DHRS9, RAI14, IDH2, MLXIP, AIP, MRPS7, HLA-DQB1*). The remaining locus *LINC00603* was identified only as genome-wide significant in the trans-ancestral meta-analysis.

A total of seven loci (three new, one independent signal, three previously reported) are concentrated in a 4 Mb segment of Chr 11 (63.2-67.2 Mb). In the previous Okada *et al.* and Kanai *et al.* East Asian and Japanese GWAS [5, 6] these loci were reported as a single locus. Köttgen *et al.* [4] reported three loci in this region (*SLC22A11, SLC22A12* and *OVOL1*). The Chr11 region is clearly of importance for serum urate control and there are more genome-wide associated loci in East Asian populations than in Europeans. At the *RELA* locus the causal variants in the East Asian population are not the same as in the European population (although we note that the Okada *et al*. [5] East Asian *RELA* signal is based entirely on imputed SNPs). Notably, the effect sizes of *SLC22A9* and *SLC22A12* (change in urate of 0.31 and 0.25 mg/dL per allele, respectively) are larger in East Asian populations than *SLC2A9* and *ABCG2* (0.18 and 0.17 mg/dL, respectively). In comparison the effect sizes in Europeans for *SLC2A9, ABCG2* and *SLC22A12* are 0.21, 0.22 and 0.07 mg/dL, respectively (note that the lead *SLC22A9* SNP *rs11231463* is uncommon in Europeans (1.1%)). In Europeans *SLC22A12* is the seventh strongest signal in serum urate after *SLC2A9, ABCG2, GCKR, SLC17A1, SLC16A9* and *INHBC*.

Very recently a separate trans-ancestral meta-analysis of the Köttgen *et al*. [4] and a new serum urate GWAS of 121,745 Japanese individuals (that encompassed all the individuals in the Kanai *et al*. [6] study) was published [38]. This study, the largest serum urate GWAS published to date, discovered 59 loci, of which 22 are newly reported beyond those reported in the Köttgen *et al*. and Kanai *et al*. studies [4, 6]. Of the 22, three overlapped with the 15 newly identified loci in the study reported here (*HLA-DQB1, B4GALT1, USP2*). Both the Kanai *et al*. [6] and Nakatochi *et al*. [38] studies reported this segment of Chr11 as a single locus. Here we dissected the Chr11 63.2-64.4 Mb segment and identified four loci in this region, including *SLC22A9* (encoding OAT7), the locus with the largest effect size on serum urate in the Japanese population (Table 1).

### Assigning causality to reported GWAS loci

We identified 44 genes with strong evidence for colocalization with a serum urate association signal (24 from the *cis*-eQTL analysis and 20 from the *trans*-eQTL analysis). Candidate causal genes at seven loci deserve brief mention (in addition to those discussed in more detail later). First, *MUC1* encodes mucin-1 (CD227), a membrane protein with excessive O-glycosylation in the extracellular domain that protects from pathogens. Mutations in *MUC1* cause autosomal dominant tubulointerstitial kidney disease [39], suggesting that regulation of this gene could influence serum urate levels via an effect on the structure and function of the kidney tubule. Second, *IGF1R* encodes the insulin-like growth factor-1 receptor, with the eQTL implicating IGF-1 signalling and resultant anabolic processes in urate control. Third, *SLC16A9* encodes mono-carboxylate transporter 9 and the urate GWAS signal is also associated with DL-carnitine and propionyl-L-carnitine levels, which are both strongly associated with serum urate levels [40]. Kidneys reabsorb carnitine from the urinary filtrate by a sodium-dependent transport mechanism [41], possibly influencing urate levels indirectly as a result of the secondary sodium dependency of urate transport [37]. Fourth, *B4GALT1* encodes *β*-1,4-galactosyltransferase 1, a Golgi apparatus membrane-bound glycoprotein. This implicates sugar modification of proteins (e.g. urate transporters) in serum urate control, either by regulating their level of expression and / or activity. Fifth, *PRPSAP1* has a *cis*-eQTL at the *QRICH2* locus - *PRPSAP1* (encoding phosphoribosyl pyrophosphate synthetase-associated protein 1) is a strong candidate gene. As a negative regulator of phosphoribosyl pyrophosphate synthetase that catalyzes the formation of phosphoribosyl pyrophosphate from ATP and ribose-5-phosphate in the purine salvage pathway decreased expression of PRPSAP1 would be predicted to contribute to increased urate levels. However, our data are not consistent with this hypothesis – *rs164009_A* associated with increased *PRPSAP1* expression and increased urate levels. Sixth, a very strong colocalized *trans*-eQTL for *CHAC2* was identified at *INHBB. CHAC2* is a y-glutamyl cyclotransferase involved in glutathione homeostasis [42, 43]. Proximal tubule cells contain high levels of glutathione which is transported in and out of the kidney via OAT1/3, MRP2/4 and OAT10 [44, 45]. Specifically, glutathione serves as a counter ion for urate reabsorption via OAT10, releasing glutathione into the lumen [45]. Thus it could be predicted that changes in *CHAC2* expression would disrupt glutathione homeostasis altering urate secretion/reabsorption in the kidney. Finally, we note that there was no evidence for a regulatory effect at *ABCG2* which is consistent with the strong evidence supporting p.Gln141Lys (*rs2231142*) as the dominant causal variant at that locus [13]. A similar scenario exists at *GCKR* where p.Leu446Pro (*rs1260326*) is the maximally-associated variant (Figure S5).

### The dystrophin complex

Of the 20 spatially supported *trans*-eQTL that colocalize with European GWAS serum urate signals, two genes, *DMD* and *UTRN* (*trans*-eQTL at *HLF* and *RREB*, respectively), also have serum urate association signals in *cis.* There was a sub-genome wide signal of association at the *UTRN* locus in the European serum urate GWAS data (*rs4896735*; *P* = 2.0 x 10^−04^) [4] and a similar signal has been reported in an Indian serum urate GWAS study (*rs12206002*; *P* < 10^−4^; not in LD with *rs4896735*) [46]. *DMD* associated with serum urate levels in the Japanese serum urate GWAS sample set (*rs1718043*; *P* = 8.8 x 10^−05^) [6]. UTRN and DMD are components of the dystrophin complex and the urate-raising alleles at these *trans*-eQTL increase expression of *UTRN* and *DMD* (Table 2). The canonical function of the dystrophin complex is well defined from its role in Duchennes Muscular Dystrophy and is crucial for stabilisation of the plasma membrane in muscle cells [47]. However syntrophins within the dystrophin complex also act as scaffolding for transporters (e.g. ABCA1 [48]) and ion channels via PDZ domains, reminiscent of the PDZK1 interaction with urate transporters [37]. Isoforms of the proteins within the dystropin complex have segment-specific distribution in the mouse nephron [49] thus it is possible that expression changes in the components of this complex in the kidney could alter the function of renal transporters that influence serum urate levels.

### OAT7

An East Asian-specific genome-wide significant signal near the gene encoding OAT7, *SLC22A9*, was confirmed. Ideally, we would have performed a colocalization analysis to assess whether this genetic association may be influencing the expression of *SLC22A9*. However, since *SLC22A9* is specifically expressed in the liver and brain and no East Asian eQTL are currently available for those tissues, this could not be performed. In lieu of providing genetic evidence that this association influences the expression of *SLC22A9*, we sought to evaluate whether OAT7 transported urate. Our data suggest that OAT7 is a very weak urate transporter in the presence of the various anions tested as exchangers (glutarate, *α*-ketoglutarate, butyrate, *β*-hydroxybutyrate). It is possible that OAT7 may function as a more efficient urate transporter in the presence of the appropriate (as yet unidentified) exchanging anion. OAT7 is a hepatic transport protein that exchanges, for the short chain fatty acid butyrate, sulphyl conjugates, xenobiotics and steroid hormones and is not inhibited by established inhibitors and substrates of other organic anion transporters such as probenecid, paraaminohippurate, nonsteroidal anti-inflammatory drugs and diuretics [36]. We found that urate transport mediated by OAT7 is inhibited by the uricosuric drugs benzbromarone and tranilast, which inhibit multiple other urate transporters [50]. Three uncommon missense variants that influence the ability of OAT7 to transport pravastatin by either causing the protein to be retained intracellularly or reducing protein levels at the plasma membrane have been reported [51], all at a frequency < 1% in East Asian. HNF4*α* plays a key role in the transactivation of the *SLC22A9* promoter [51], an interesting observation given that HNF4*α* is also required for expression of the gene encoding the urate transportosome-stabilizing molecule PDZK1 in the liver [11], and is implicated in control of serum urate levels via the *MAFTRR* locus [12].

### Colocalization analysis assigns causation to variants at MLXIPL and MLXIP

We identified the paralogs *MLXIPL* and *MLXIP* as the putative causal genes at the *BAZ1B* and *MLXIP* loci respectively. These genes encode the ChREBP and MondoA proteins, which are glucose-sensitive transcription factors involved in energy metabolism – including glycolytic targets and glycolysis [52–54]. These proteins form heterodimers with the Mlx protein, and both of these proteins are activated by high levels of intracellular glucose-6-phosphate – a product of the first step of the glycolysis and pentose phosphate pathways. Increased activity of the pentose phosphate pathway leads to the production of ribose-5-phosphate thus stimulating *de novo* purine nucleotide synthesis. The resulting nucleotides are ultimately catabolised into urate if they are not otherwise utilized. In *Drosophila* at least, the ChREBP/Mondo-Mlx complex is responsible for the majority of transcriptional changes that result from glucose consumption, including the pentose phosphate pathway [52]. The colocalization results reveal that the serum urate-increasing variants at both loci decrease expression of *MLXIPL* and *MLXIP*. Taken together, this suggests a possible mechanism whereby the decreased basal expression of ChREBP snd MondoA results in increased activity of the pentose phosphate pathway and therefore higher levels of serum urate.

### MRPS7 and IDH2 and mitochondrial function

*MRPS7* is putatively involved in serum urate control via mitochondrial processes. Of relevance, reduced relative mitochondrial DNA copy number is associated with gout [55]. The association signal at the *MRPS7* locus colocalized with gene expression of *MRPS7* and *GGA3. MRPS7* encodes the mitochondrial ribosomal protein S7, which is required for the assembly of the small ribosomal subunit of the mitochondria. A whole exome study revealed that a non-synonymous mutation in *MRPS7* (p.Met184Val), which destabilizes the protein and reduces expression, results in impaired mitochondrial protein synthesis and impaired mitochondrial function [56]. The patients in this study presented with congenital sensorineural and significant hepatic and renal impairment, consistent with a role for reduced MRPS7 activity in renal function. Our findings show that the urate-increasing G-allele decreases the expression of *MRPS7* (Table 2), consistent with the hypothesis generated by the p.Met184Val phenotype. Also implicating mitochondrial function is *IDH2* which encodes isocitrate dehydrogenase that catalyzes the decarboxylation of isocitrate to 2-oxyglutarate in the citric acid cycle. The urate-increasing allele associates with reduced expression of *IDH2*. Somatic mutations in *IDH2* are implicated in a range of diseases including cancers such as glioma and acute myeloid leukemia (where an inhibitor is in phase III clinical trial [57]) and the tumor syndromes Ollier disease and Maffucci syndrome [58]. Understanding the molecular mechanism of urate control by the *MRPS7* and *IDH2* loci locus could lead to insights into the mitochondrial processes that influence serum urate levels.

### Trans-ancestral functional fine-mapping identifies putative causal variants

To connect GWAS loci where we identified candidate causal genes to an underlying causal variant, we performed trans-ancestral fine-mapping with PAINTOR using the kidney, gastrointestinal tract, and liver cell type group annotations as functional priors. We identified six loci (*RREB1, INHBC, HLF, UBE2Q2, SFMBT1, HNF4G*) that had colocalized eQTL and contained SNPs with high posterior probabilities of causality (>0.8). Two additional loci *SLC2A9* and *SLC22A12* also contained SNPs with high posterior probabilities of causality (>0.8) that were *cis* and *trans*-eQTL for *RP11-448G15.1* and *RNF169,* respectively. Many of these SNPs overlapped annotated regulatory regions of the genome (Table 3). These candidate causal variants and genes provide a starting point for understanding how these variants alter serum urate levels. The power of this approach is illustrated in our prior work on the *PDZK1* locus [11]. Here, we experimentally confirmed that *PDZK1* was the causal gene, with *rs1967017* (one of the two candidate causal variants identified with posterior probabilities >0.25 (Table S1)) being a highly likely causal variant via altering a binding site for hepatocyte nuclear factor 4*α*. We have also applied a similar approach to the *MAF* locus [12]. *MAF* is a complex locus with population-specific signals, and for one of these signals we experimentally demonstrated that the effect on urate arises from one of two SNPs within a kidney specific enhancer that is co-expressed with *MAF* and *HNF4A* in the developing proximal tubule. This study also identified colocalised eQTL for two long intergenic non-coding RNAs *MAFTRR* and *LINC01229* that regulate *MAF* expression in *cis*, and other genes implicated in urate metabolism in *trans* [12]. These studies highlight the power of initially combining colocalization analyses and fine-mapping using prior information to determine the molecular mechanisms that underlie GWAS signals.

In conclusion, we have identified 15 new GWAS signals associated with serum urate levels. By *cis*-eQTL colocalization we identified 24 candidate causal genes and by *trans*-eQTL analysis we implicated a further 20 genes in the molecular control of serum urate levels. Highlighted insights into molecular mechanisms come from identification of the protein encoded by *SLC22A9* (OAT7) to be a urate transporter, the implication of mitochondrial function via *MRPS7*, the identification of *MLXIP* (alongside the already identified *MLXIPL*) and intriguing data genetically implicating the dystrophin complex in control of serum urate levels.

## Methods

### Data preparation and quality control

Summary statistics from the Global Urate Genetics Consortium (GUGC) meta-analysis of GWAS data consisting of 110,238 individuals of European ancestry [4] (http://metabolomics.helmholtz-muenchen.de/gugc/), and a meta-analysis consisting of 21,417 individuals of East Asian ancestry [5] were utilised. For both datasets the following quality control procedure was followed. Firstly, we removed any SNPs that were not present in the Phase 3 release of the 1000 Genomes for the representative populations (EUR and EAS), or where the alleles were not identical between this summary data and the 1000 Genomes (e.g. the alleles were G/T in the GUGC meta-analysis and T/A in the 1000 Genomes dataset) [59]. The effective sample size for each SNP was calculated using the Genome-wide Complex Trait Analysis (GCTA, v1.25.2) toolkit [60] and SNPs with effective sample sizes > 2 standard deviations from the mean were excluded. Finally, SNPs with a minor allele frequency (MAF) of less than 0.01 were excluded.

### Trans-ancestral meta-analysis

ImpG (v1.0) was used to impute Z-scores into the European and East Asian summary statistics. For the reference haplotypes the Phase 3 release of the 1000 Genomes project was used [59], and only bi-allelic SNP markers having a minor allele frequency greater than 0.01 in the relevant population were included. All imputed markers with a predicted R^2^ of less than 0.8 were removed. Meta-analysis was performed by summing the Z-scores and weighting by sample size. For the imputed SNPs, the sample size was estimated as the median of the sample size of the SNPs where this information was available. To provide an adjustment for inflated test statistics, the LD-score intercept in the original summary statistics files was calculated using LD-score regression [61]. This intercept adjusts the test statistics for confounding, such as cryptic relatedness, but in contrast to genomic control will not remove inflation caused by a true polygenic signal.

Independent regions were identified using the following protocol. Firstly, SNPs that were genome-wide significant (*P* < 5 x 10^−08^) were padded 50 kb either side of the SNP position, and all overlapping regions were clumped together. Secondly, the maximal R^2^ > 0.6 for the most significant SNP in each of these regions was calculated for each population. Finally, the maximal regions from the P-value clumping and LD approach were created, and any overlapping regions were merged. SNPs that were not present in both datasets were also analyzed, and for those SNPs the LD was only calculated in the relevant population. Based on their proximity to stronger signals four loci (*Chr4/rs114188639/CLNK1, Chr8/rs2927238/HNF4G, Chr11/rs641811/FLRT1, Chr11/rs117595559/VPS51*) were visually examined by LocusZoom and subjected to conditional analysis – of these only *rs641811 / FLRT1* was concluded to be independent of the nearby signal. For all significant SNPs, the meta-analysis effect estimate was calculated using the inverse variance method, and when there was no effect estimate, it was estimated from the Z score using the following equation (1).

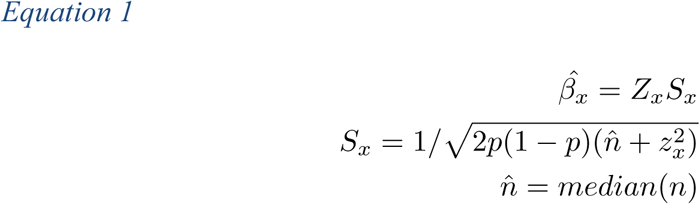

### Conditional analysis

A conditional and joint analysis of the European summary statistics for all genome-wide significant regions identified by meta-analysis was done. This was not done on the East Asian summary statistics owing to the lack of availability of both a LD matrix and a reference haplotype set of sufficient size. For all imputed SNPs the effect estimate was calculated as above (equation 1). The genotypic data from the UK Biobank was used as the reference for the LD, and to improve computational efficiency only a random 15% (22,872) of samples were included. Since the GCTA-COJO module [62] was not designed to utilize dosage matrices, we performed this analysis using our own software, Correlation-based Conditional analysis (COCO: https://github.com/theboocock/coco) which, based on the methods presented in GCTA-COJO [62], was designed to perform conditional and joint analysis from summary statistics with some minor alterations to use LD correlation matrices as input. To discover conditional associations, the coco pipeline implemented a forward stepwise selection using a residual-based regression. First, SNPs were ranked on marginal test statistics, then the top SNP was selected and the result of extracting the residuals from this model and performing a regression with every other SNP was estimated. These test-statistics were then ranked. If the new top SNP passed the P-value threshold it was added to a joint model with the other selected SNP, which was used as the new model for residual extraction. This process was then repeated until no SNPs passed the significance threshold. In practice, we restricted the maximum number of selected SNPs at a locus to five (there is evidence for multiple signals at *SLC2A9* [10]), and we did not consider any pairs of SNPs having an R^2^ > 0.9. To ensure that the method was working correctly, simple phenotypes were simulated and it was verified that COCO yields almost identical results to the lm function in the R programming language.

A mathematical explanation of the method is given as follows. We assume we have mean centered genotypes in a matrix X. To perform GWAS we generate a marginal statistic for each variant individually (Equation 2).

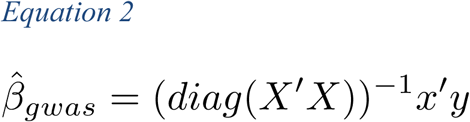

Using substitution into the ordinary least squares equations we can convert these marginal effects into joint effects, and also calculate the standard error (Equation 3).

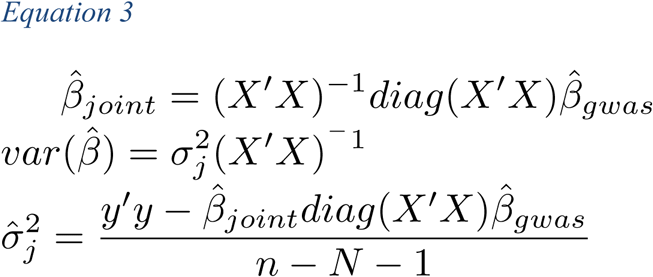

Where N is equal to the number of SNPs in the joint model. Finally, we can approximate a regression of the residuals from a joint model (Equation 4).

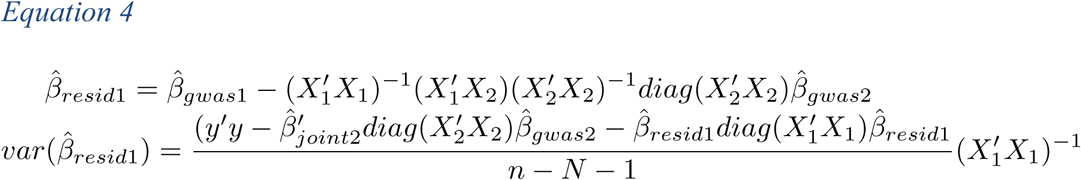

Where X_1_ is the genotype matrix of SNPs to be regressed on by the residuals, X_2_ is the genotype matrix of the joint model, and N is equal to the number of SNPs in the residual model. In practice the data matrix X is unavailable as summary statistics were used, but it is possible to approximate this matrix using the LD structure from a reference panel (Equation 5).

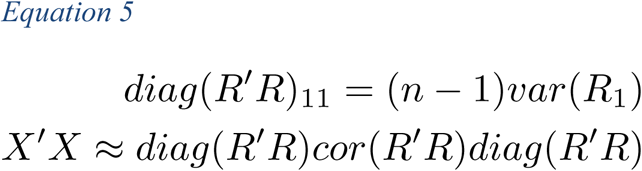

Where R is the reference genotype matrix, sigma is the LD matrix for the locus, and the diagonal of R*′*R is modified to be equal to the sample size of the SNP minus one in the GWAS multiplied by the genotypic variance of the SNP observed in the reference panel. Since the data were generated from dosages and not hard-called genotypes, using the observed genotypic variance in the reference panel would have accounted for some of the uncertainty introduced by imputation.

To calculate the effective number of hypothesis tests, Eigen value decomposition was performed on the SNP correlation matrix for each region, using data from the European individuals from the 1000 Genomes Project. The number of hypotheses tested per region was calculated as the number of Eigen values that were required to explain 0.995 of the total sum of the Eigen values. The total number of hypotheses tested in the conditional analysis was taken as the sum of the per region hypothesis counts [63]. This revealed that in the focused conditional analysis, we were performing approximately 5,443 hypothesis tests. The multiple-testing threshold for our conditional analysis was therefore determined to be 9.2 x 10^−06^ (0.05/5443).

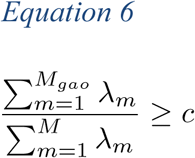

The following equation was used to calculate variance explained by each SNP in the meta-analysis and joint analysis (Equation 7).

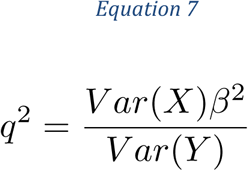

Where Var(X) was the variance for each SNP, calculated as 2p(1 - p) with p as the allele frequency. *β* was the effect estimate, and Var(Y) was calculated as the pooled variance estimate provided by the GCTA software when performing the conditional analysis. This pooled variance was calculated using the equation below (Equation 8).

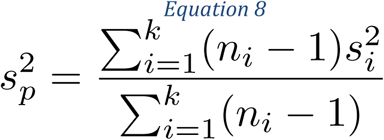

Where *s_i_^2^* was the phenotypic variance estimated by the GCTA software for each chromosome, and *n_i_* was the number of SNPs on each chromosome. This revealed that the empirical variance of sex adjusted serum urate was 1.624. Unadjusted variance in serum urate was also calculated using the equation above, where *s_i_^2^* was replaced with the variance in serum urate for each study, and *n_i_* was replaced with the number of participants in each study. This analysis revealed that the empirical variance of unadjusted serum urate was 1.964.

### Heritability and functional enrichments

LD Score regression was used to partition SNP heritability of serum urate [61]. An estimate was generated using LD Score for the amount of heritability explained by all SNPs additively in the Köttgen *et al*. [4] meta-analysis. We also performed functional annotation-partitioned LD score regression to determine which cell type groups and cell types contribute significantly to the heritability of serum urate [64]. The comprehensive set of functional annotations that were released with partitioned LD Score regression (https://data.broadinstitute.org/alkesgroup/LDSCORE/) were used. This works by comparing the results to a baseline model that contains annotations such as evolutionary conservation, and pooled cell type annotations such as DNase1 hypersensitivity. We calculated P-values and Bonferonni-corrected thresholds by dividing by the total number of tests within each of the cell type group and cell type specific analyses, noting that this is a conservative adjustment because the annotations are correlated. Benjamini-Hochberg false discovery rate (FDR) adjusted P-values were also calculated [65]. All results were visualized using ggplot2 [66].

### Functional trans-ancestral fine mapping with PAINTOR

PAINTOR (v3.0) [67] was initially used to fine map the 38 loci associated at a genome-wide level of significance in this study with serum urate within the separate European and East Asian GWAS. This initial analysis revealed that both the *RELA* and *HLA* loci were inappropriate loci for trans-ancestral fine-mapping. For the *RELA* locus, the association signal between the European and East Asian GWAS clearly involves different causal variants. For *HLA-DQB1*, the large number of SNPs in the region resulted in computational errors in the PAINTOR software. Both of the loci were excluded from all additional PAINTOR analyses. Cell type groups that were significant in the LD score regression analysis were used with PAINTOR. To assess how much the East Asian GWAS and these functional annotations improved serum urate fine-mapping, the average size of the 90% causal credible sets in three analyses was calculated.

1. European GWAS.
2. European and East Asian GWAS.
3. European and East Asian GWAS and functional annotations.

### Cis-eQTL identification

We used COLOC [68] to colocalize the urate-associated loci with publicly available eQTL data from the Genotype Tissue Expression Project (GTEx v6p). COLOC is a Bayesian method that compares four different statistical models at a locus. These models are: no causal variant in the GWAS or the eQTL region; a causal variant in either the GWAS or the eQTL region, but not both; different causal variants in the GWAS and the eQTL region; or a shared causal variant in the GWAS and the eQTL region. All the *cis*-eQTL regions from a GTEx tissue were merged with the genome-wide European serum urate GWAS data. Genes that were annotated as novel transcripts were removed. For learning the priors each *cis*-eQTL region was treated as independent and the likelihood was maximized using the Nelder-mead algorithm. Genes that had a posterior probability of colocalization greater than 0.8 were considered to have a shared causal variant with serum urate. We did not restrict our analysis only to the genome-wide significant loci, which made it possible to identify novel serum urate loci. If multiple tissues supported colocalization at probability > 0.8 the posterior probability was averaged.

### Trans-eQTL identification

The Contextualize Developmental SNPs using 3D Information (CoDeS3D) algorithm (GitHub, https://github.com/alcamerone/codes3d) [69] was used to identify long-distance regulatory relationships for serum urate-associated SNPs. This analysis leverages known spatial associations from Hi-C databases [70] and gene expression associations (eQTL data from the GTEx catalogue [71]) to assess regulatory connections. Briefly, SNPs were mapped onto Hi-C restriction fragments, the genes that physically interact with these restriction fragments identified and collated (SNP-gene spatial pairs). SNP-gene pairs were screened through GTEx to identify eQTL. The FDR was calculated using a stepwise Benjamini-Hochberg correction procedure and incorporated the number of tests and eQTL value list. An FDR value of < 0.05 was accepted as statistically significant [69]. COLOC was then used to co-localize *trans*-eQTL with serum urate GWAS signals.

### Gout case-control sample sets for replicating serum urate associations

The Japanese gout data set, generated as previously described, [30] consisted of 945 male gout patients and 1,213 male controls, where gout was clinically ascertained. The Chinese data set, generated as previously described [32], consisted of 1,255 male gout cases and 1,848 male control where gout was clinically ascertained according to the American College of Rheumatology diagnostic criteria. The European gout dataset was generated from 7,342 gout patients and 352,534 controls of European ancestry from the UK Biobank [31], where gout was ascertained by self-report of physician-diagnosed gout or use of urate-lowering therapy [72]. Gout association in UK Biobank was tested using logistic regression, adjusted by age, sex, and the first 10 principal components (out of 40).

### SLC22A9 – Cell lines, RNA Extraction and RT-PCR

Human kidney proximal tubule epithelial cell line (PTC-05) was obtained from Ulrich Hopfer (Case Western Reserve University, Cleveland, Ohio) and grown (37°C in a 5% CO_2_) on type IV collagen-coated Petri dish in a 1:1 mixture of DMEM and HAM’S F12 media containing 5 mM glucose, 10% fetal bovine serum (FBS), 2 mM glutamine, 1 mM pyruvate, 5 µg/ml transferrin, 5 µg/ml insulin, 10 ng/ml human epidermal growth factor, 4 µg/ml dexamethasone, 15 mM HEPES (pH 7. 4), 0.06% NaHCO_3_, 10 ng/ml interferon-gamma, 50 µM ascorbic acid, 20 nM sodium selenite (Na_2_SeO_3_), 1nM triiodothyronin (T3) and penicillin (50 units/ml) / streptomycin (50 µg/ml). Human embryonic kidney HEK293 cells (ATCC) and human hepatocellular carcinoma HEPG2 cells (ATCC) were grown (37°C in a 5% CO_2_) and maintained in Dulbecco’s Modified Eagle’s Medium (DMEM) and Eagle’s Minimum Essential Media (EMEM), respectively, supplemented with 4.5 g/L glucose, 2 mM glutamine, 1 mM sodium pyruvate, 10% FBS and penicillin (50 units/ml)/streptomycin (50 µg/ml).

Total RNA from human cell lines (PTC-05, HepG2 and HEK-293T) was extracted using spin columns with the RNeasy Mini Kit (QIAGEN, GmbH, Germany) following the manufacturer’s instructions. Approximately 2 μg of DNase-treated total RNA, isolated from cells, was primed with poly-dT and random hexamers and then reverse-transcribed using AMV reverse transcriptase (New England Biolabs, Ipswich, MA). An equal amount of cDNA was used for PCR amplification of OAT7 and GAPDH cDNAs using the following primers, followed by electrophoresis.

hOAT7-2S [sense] 5′-CAACCTCAATGGCCTTTCAGGACCTCCTGG-3′ hOAT7-3A [antisense] 5′-GCCTGGAATCTGTGTGTTGCCCACTCGG-3′ hGAPDH-1S [sense] 5′-CGGAGTCAACGGATTTGGTCGTATTG-3′ hGAPDH-1A [antisense] 5′-GACTGTGGTCATGAGTCCTTCCACGA-3′

### SLC22A9 - urate transport analysis of OAT7

Studies using *Xenopus laevis* oocytes were performed in accordance with the Guide for the Care and Use of Laboratory Animals as adopted and promulgated by the U.S. National Institutes of Health, and were approved by the Institution’s Animal Care and Use Committee. Mature female *Xenopus laevis* frogs (NASCO, Fort Atkinson, MI) were subjected to partial ovariectomy under tricane (SIGMA St Louis, MO) anesthesia (0.17% for 15–20 min) as described previously [50]. A small incision was made in the abdomen and a lobe of ovary was removed. Subsequently, the oocytes were pre-washed for 20 min in Ca^2+^-free ND96 medium (96 mM NaCl, 2 mM KCl, 1 mM MgCl_2_, 5 mM HEPES, pH 7.4) to remove blood and damaged tissue. Oocytes were then defolliculated by treatment with 3.5 mg/ml of collagenase enzyme (Roche, Indianapolis, IN) in Ca^2+^-free ND96 medium for about 120 min with gentle agitation at room temperature (25°C). Subsequent to this treatment, oocytes were washed three times with ND96 medium, and incubated (16-18 °C) in isotonic Ca^2+^-containing ND96 medium (96 mM NaCl, 2.0 mM KCl, 1.8 mM CaCl_2_, 1.0 mM MgCl_2_ and 5 mM Hepes, pH 7.4) supplemented with 2.5 mM pyruvate and gentamycin (10 µg/ml).

For expression of OAT7 and OAT1 in *Xenopus laevis* oocytes, their respective full-length cDNAs were cloned into the pGEMHE vector, wherein the cDNA insert is flanked by the *Xenopus laevis β*-globin 5′-UTR and 3′-UTR [73]. These constructs were linearized and cRNAs were synthesized *in vitro* using T7 RNA polymerase (mMESSAGE mMACHINE; Ambion, Austin, TX) following the supplier’s protocol. Isopropanol-precipitated, *in vitro* transcribed capped cRNAs were washed twice with 70% ethanol, the cRNA pellet was dried and then dissolved in sterile nuclease-free water. The yield and integrity of the capped cRNA samples was assessed by spectroscopy (at 260 nm) and 1% agarose-formaldehyde gel electrophoresis respectively. All cRNA samples were stored frozen in aliquots at -80 °C until used.

About 18 hours after isolation, oocytes were microinjected with 50 nl of sterile water, 50 mM tris pH 7.4, or 50 nl of a cRNA solution in 50 mM tris buffer (pH 7.4) containing 25 ng of the indicated cRNA using fine-tipped micropipettes by a microinjector (World Precision Instrument Inc. Sarasota, FL). The microinjected oocytes were then incubated in isotonic ND96 medium (pH 7.4) containing 1.8 mM CaCl_2_, 2.5 mM pyruvate, gentamycin (10µg/ml) at 16-18 °C for approximately 48 h to allow expression of protein from microinjected cRNA.

For [^14^C]-urate (specific activity: 50 mCi/mmol) uptake experiments in *Xenopus laevis* oocytes, oocytes expressing proteins as indicated (OAT7 and OAT1) were washed four times with ND96 medium (96 mM NaCl, 2.0 mM KCl, 1.8 mM CaCl_2_, 1.0 mM MgCl_2_ and 5 mM Hepes, pH 7.4) without pyruvate and gentamycin. OAT7 functions as a butyrate exchanger [36], therefore OAT7-expressing oocytes were microinjected with 50 nl of 100 mM butyrate to optimize urate transport by “trans-activation” [50]. After approximately 60 min of starvation, oocytes were preincubated in the ND96 uptake medium for 30 min before incubation (25°C, in a horizontal shaker-incubator) in the uptake medium containing [^14^C]-urate (40 µM). After 60 min of incubation in the uptake medium, oocytes (20 per group) were washed three times with ice-cold uptake medium to remove external adhering radioisotope. OAT7-expressing oocytes were then exposed to DMSO (diluent for uricosurics) or the uricosuric drugs tranilast and benzbromarone, as indicated. The radioisotope content of each individual oocyte was measured by scintillation counter following solubilization in 0.3 ml of 10% (v/v) SDS and addition of 2.5 ml of scintillation fluid (Ecoscint). All uptake experiments included at least 20 oocytes in each experimental group; statistical significance was defined as two-tailed *P* < 0.05, and results were reported as means ± S. E. Statistical analyses including linear regressions and significance were determined by Student’s t test using SigmaPlot software.

## Acknowledgements

This research has been conducted using the UK Biobank Resource under Application Number 12611. Akiyoshi Nakayama is thanked for statistical analytic support. The Health Research Council of New Zealand is acknowledged for funding support.

## Supporting Information

**Supplementary Figure 1. Q-Q plots for serum urate.**

Quantile-quantile plot showing observed P-values versus expected P-values. Q-Q curves are provided for: all SNPs, excluding highly significant loci (*P*<1E-20), and excluding GWAS significant loci (*P*<5E-08). Genomic-control is provided with and without the ImpG imputed SNPs. The LD-score intercept is provided for the European and East Asian GWAS.

**Supplementary Figure 2. Regional associations plot of the 3 undescribed serum urate loci identified in the Okada *et al.* East Asian GWAS.**

Regional association plots of the 3 previously undescribed Chr11 serum urate loci (*SLC22A9, PLA2G16* and *AIP*) that were identified in the East Asian GWAS. The lead SNPs are indicated by a purple dot. The color of the surrounding SNPs indicates the strength of LD with the lead SNP according to the key in the left top hand corner, measured as *r^2^* found in the HapMap data (hg19/1000 genomes Nov 2014) East Asian. The plots were generated using LocusZoom.

**Supplementary Figure 3. Regional association plots for the 7 novel serum urate loci identified in the trans-ancestral meta-analysis.**

Regional association plots of the 7 novel serum urate loci (*FGF, LINC00603, HLA-DQB1, B4GALT1, BICC1, FLRT1* and *USP2*) that were identified by trans-ancestral meta-analysis. The lead SNPs are indicated by a purple dot. The color of the surrounding SNPs indicates the strength of LD with the lead SNP according to the key in the left top hand corner, measured as *r^2^* found in the HapMap data (hg19/1000 genomes Nov 2014). European LD data were utilized as the reference for the trans-ancestral regional association plots. The plots were generated using LocusZoom.

**Supplementary Figure 4. Regional association plot of the *RELA* locus reveals an East Asian specific association with serum urate levels.**

Regional association plots at the *RELA* locus for both the European and East Asian GWAS are shown. The lead SNPs are indicated by a purple dot. The color of the surrounding SNPs indicates the strength of LD with the lead SNP according to the key in the left top hand corner, measured as *r^2^* found in the HapMap data (hg19/1000 genomes Nov 2014). a) Regional association plot with LD calculated from the index European SNP variant *rs12289836*. b) Regional association plot with LD calculated from the East Asian specific variant *rs1227200*. The plots were generated using LocusZoom.

**Supplementary Figure 5. Regional association plots of all significant (*P* < 5E-08) serum urate loci.**

For each locus, we have provided regional association plots for the East Asian and European GWAS in addition to the trans-ancestral GWAS. The lead SNPs are indicated by a purple dot. The color of the surrounding SNPs indicates the strength of LD with the lead SNP according to the key in the left top hand corner, measured as *r^2^* found in the HapMap data (hg19/1000 genomes Nov 2014). European LD data were utilized as the reference for the trans-ancestral regional association plots. The plots were generated using LocusZoom.

**Supplementary Figure 6. Regional association plot of the *MAF* locus reveals an East Asian-specific association and a shared association with serum urate levels.**

Regional association plots at the *MAF* locus from both the European and East Asian serum urate GWAS are shown. The lead SNPs are indicated by a purple dot. The color of the surrounding SNPs indicates the strength of LD with the lead SNP according to the key in the left top hand corner, measured as *r^2^* found in the HapMap data (hg19/1000 genomes Nov 2014). (A) Regional association plot with LD calculated from the shared trans-ancestral variant *rs1150189*. (B) Regional association plot with LD calculated from the East Asian-specific variant *rs889472*. The plots were generated using LocusZoom.

**Supplementary Figure 7. Regional association plots of eQTL colocalization.**

For each locus, regional association plots for the serum urate locus (European GWAS) and the GTEx eQTL results are shown. For any gene that colocalized with a serum urate locus in multiple-tissues, we only show one representative figure pair. For serum urate loci with multiple colocalized genes, we show one representative figure pair for each colocalized gene. The lead SNPs are indicated by a purple dot. The color of the surrounding SNPs indicates the strength of LD with the lead SNP according to the key in the left top hand corner, measured as *r^2^* found in the European HapMap data (hg19/1000 genomes Nov 2014). The plots were generated using LocusZoom.

**Supplementary Figure 8. Regional association plots of the *DMD* and *UTRN* loci.**

Regional association plots at *UTRN* (LD calculated from lead SNP rs4896735 using European LD from 1000 genomes 2014) from the European urate GWAS and *DMD* from the Japanese urate GWAS (LD calculated from lead SNP rs171843 using Asian LD from 1000 genomes 2014) are shown. The plot was generated using LocusZoom.

**Supplementary Figure 9. Cell type-specific functional heritability enrichments.**

Cell type-specific enrichments for serum urate levels. The color of each bar represents the cell type group of each annotation. The direction of enrichment is indicated by adding a sign to the –log10 P-value. A positive sign indicates enrichment and a negative sign indicates depletion. These results are displayed in four panels one for each histone mark: a) H3K27ac ChIP-seq; b) H3K9ac; c) H3K4me3; d) H3K4me1.

**Supplementary Figure 10. Regional association plots of *RP11-448G15.1* and *RNF169* eQTL.** For each locus, regional association plots for the serum urate locus (European GWAS) and the GTEx eQTL results are shown. The lead SNPs are indicated by a purple dot. The color of the surrounding SNPs indicates the strength of LD with the lead SNP according to the key in the left top hand corner, measured as *r^2^* found in the European HapMap data (hg19/1000 genomes Nov 2014). The plots were generated using LocusZoom.

**Supplementary Table 1. Partitioned serum urate heritability enrichment estimates for cell-type groups.** These enrichments were generated using LD-score functional partitioning of the European GWAS summary statistics.

**Supplementary Table 2. Partitioned serum urate heritability enrichment estimates for cell-type specific epigenomic profiles.** These enrichments were generated using LD-score functional partitioning of the European GWAS summary statistics.

**Supplementary Table 3. PAINTOR results.** Trans-ancestral functional fine-mapping results for 36 serum urate loci (excluding *MHC* and *RELA*). Loci can be distinguished by their index SNP and locus names. The results are summarized from three PAINTOR models: the model which used the GWAS data from both the East Asian and European population, the model which used the GWAS data from the East Asian population only, and the model which used the GWAS data from the European population only. All models included significant cell type group annotations.

**Supplementary Table 4. Functionally annotated variants identified by PAINTOR.** Haploreg v4.1 was used to annotate all variants with a PAINTOR posterior probability > 0.8. Annotations include enhancer histone marks, DNAse peaks, proteins bound (ChIP), motifs changed by the alternate allele, number of GWAS and eQTL hits and location with respect to the nearest gene.

## Notes

https://zenodo.org/record/3366490#.XVJbLZPYryI

## References

1. Kuo CF, Grainge MJ, Zhang W, Doherty M. Global epidemiology of gout: prevalence, incidence and risk factors. Nat Rev Rheumatol. 2015;11(11):649–62.

2. Dalbeth N, Merriman TR, Stamp LK. Gout. The Lancet. 2016;388(10055):2039–52.

3. Martinon F, Petrilli V, Mayor A, Tardivel A, Tschopp J. Gout-associated uric acid crystals activate the NALP3 inflammasome. Nature. 2006;440(7081):237–41.

4. Kottgen A, Albrecht E, Teumer A, Vitart V, Krumsiek J, Hundertmark C, et al. Genome-wide association analyses identify 18 new loci associated with serum urate concentrations. Nat Genet. 2013;45(2):145–54.

5. Okada Y, Sim X, Go MJ, Wu JY, Gu D, Takeuchi F, et al. Meta-analysis identifies multiple loci associated with kidney function-related traits in east Asian populations. Nat Genet. 2012;44(8):904–9.

6. Kanai M, Akiyama M, Takahashi A, Matoba N, Momozawa Y, Ikeda M, et al. Genetic analysis of quantitative traits in the Japanese population links cell types to complex human diseases. Nat Genet. 2018;50(3):390–400.

7. Phipps-Green AJ, Merriman ME, Topless R, Altaf S, Montgomery GW, Franklin C, et al. Twenty-eight loci that influence serum urate levels: analysis of association with gout. Ann Rheum Dis. 2016;75(1):124–30.

8. Urano W, Taniguchi A, Inoue E, Sekita C, Ichikawa N, Koseki Y, et al. Effect of genetic polymorphisms on development of gout. J Rheumatol. 2013;40(8):1374–8.

9. Major TJ, Dalbeth N, Stahl EA, Merriman TR. An update on the genetics of hyperuricaemia and gout. Nat Rev Rheumatol. 2018;14(6):341–53.

10. Merriman TR. An update on the genetic architecture of hyperuricemia and gout. Arthritis Res Ther. 2015;17(1):98.

11. Ketharnathan S, Leask M, Boocock J, Phipps-Green AJ, Antony J, O’Sullivan JM, et al. A non-coding genetic variant maximally associated with serum urate levels is functionally linked to HNF4A-dependent PDZK1 expression. Hum Mol Genet. 2018;27(22):3964–73.

12. Leask M, Dowdle A, Salvesen H, Topless R, Fadason T, Wei W, et al. Functional Urate-Associated Genetic Variants Influence Expression of lincRNAs LINC01229 and MAFTRR. Front Genet. 2018; DOI: 10.3389/fgene.2018.00733.

13. Cleophas MC, Joosten LA, Stamp LK, Dalbeth N, Woodward OM, Merriman TR. ABCG2 polymorphisms in gout: insights into disease susceptibility and treatment approaches. Pharmgenomics Pers Med. 2017;10:129–42.

14. Ichida K, Matsuo H, Takada T, Nakayama A, Murakami K, Shimizu T, et al. Decreased extra-renal urate excretion is a common cause of hyperuricemia. Nat Commun. 2012;3:764.

15. Matsuo H, Takada T, Ichida K, Nakamura T, Nakayama A, Ikebuchi Y, et al. Common defects of ABCG2, a high-capacity urate exporter, cause gout: a function-based genetic analysis in a Japanese population. Sci Transl Med. 2009;1(5):5ra11.

16. Nakayama A, Matsuo H, Nakaoka H, Nakamura T, Nakashima H, Takada Y, et al. Common dysfunctional variants of ABCG2 have stronger impact on hyperuricemia progression than typical environmental risk factors. Sci Rep. 2014;4:5227.

17. Woodward OM, Tukaye DN, Cui J, Greenwell P, Constantoulakis LM, Parker BS, et al. Gout-causing Q141K mutation in ABCG2 leads to instability of the nucleotide-binding domain and can be corrected with small molecules. Proc Natl Acad Sci U S A. 2013;110(13):5223–8.

18. Morris AP. Transethnic meta-analysis of genomewide association studies. Genet Epidemiol. 2011;35(8):809–22.

19. Zaitlen N, Pasaniuc B, Gur T, Ziv E, Halperin E. Leveraging genetic variability across populations for the identification of causal variants. Am J Hum Genet. 2010;86(1):23–33.

20. Consortium EP. An integrated encyclopedia of DNA elements in the human genome. Nature. 2012;489(7414):57.

21. Roadmap Epigenomics C, Kundaje A, Meuleman W, Ernst J, Bilenky M, Yen A, et al. Integrative analysis of 111 reference human epigenomes. Nature. 2015;518(7539):317–30.

22. Lonsdale J, Thomas J, Salvatore M, Phillips R, Lo E, Shad S, et al. The Genotype-Tissue Expression (GTEx) project. Nat Genet. 2013;45(6):580–5.

23. Schierding W, Antony J, Cutfield WS, Horsfield JA, O’Sullivan JM. Intergenic GWAS SNPs are key components of the spatial and regulatory network for human growth. Hum Mol Genet. 2016;25(15):3372–82.

24. Fadason T, Schierding W, Lumley T, O’Sullivan JM. Chromatin interactions and expression quantitative trait loci reveal genetic drivers of multimorbidities. Nat Commun. 2018;9(1):5198.

25. Võsa U, Claringbould A, Westra H-J, Bonder MJ, Deelen P, Zeng B, et al. Unraveling the polygenic architecture of complex traits using blood eQTL meta-analysis. bioRxiv. 2018:447367.

26. Cruz-Tapias P, Perez-Fernandez OM, Rojas-Villarraga A, Rodriguez-Rodriguez A, Arango MT, Anaya JM. Shared HLA Class II in Six Autoimmune Diseases in Latin America: A Meta-Analysis. Autoimmune Dis. 2012;DOI: 10.1155/2012/569728.

27. Dobbyn A, Huckins LM, Boocock J, Sloofman LG, Glicksberg BS, Giambartolomei C, et al. Co-localization of Conditional eQTL and GWAS Signatures in Schizophrenia. bioRxiv. 2017:129429.

28. Wen CC, Yee SW, Liang X, Hoffmann TJ, Kvale MN, Banda Y, et al. Genome-wide association study identifies ABCG2 (BCRP) as an allopurinol transporter and a determinant of drug response. Clin Pharmacol Ther. 2015;97(5):518–25.

29. Kamatani Y, Matsuda K, Okada Y, Kubo M, Hosono N, Daigo Y, et al. Genome-wide association study of hematological and biochemical traits in a Japanese population. Nat Genet. 2010;42(3):210–5.

30. Matsuo H, Yamamoto K, Nakaoka H, Nakayama A, Sakiyama M, Chiba T, et al. Genome-wide association study of clinically defined gout identifies multiple risk loci and its association with clinical subtypes. Ann Rheum Dis. 2016;75(4):652–9.

31. Bycroft C, Freeman C, Petkova D, Band G, Elliott LT, Sharp K, et al. The UK Biobank resource with deep phenotyping and genomic data. Nature. 2018;562(7726):203–9.

32. Li C, Li Z, Liu S, Wang C, Han L, Cui L, et al. Genome-wide association analysis identifies three new risk loci for gout arthritis in Han Chinese. Nat Commun. 2015;6:7041.

33. Scharpf RB, Mireles L, Yang Q, Kottgen A, Ruczinski I, Susztak K, et al. Copy number polymorphisms near SLC2A9 are associated with serum uric acid concentrations. BMC Genet. 2014;15(1):81.

34. Wei WH, Guo Y, Kindt AS, Merriman TR, Semple CA, Wang K, et al. Abundant local interactions in the 4p16.1 region suggest functional mechanisms underlying SLC2A9 associations with human serum uric acid. Hum Mol Genet. 2014;23(19):5061–8.

35. Kheradpour P, Kellis M. Systematic discovery and characterization of regulatory motifs in ENCODE TF binding experiments. Nucleic Acids Res. 2014;42(5):2976–87.

36. Shin HJ, Anzai N, Enomoto A, He X, Kim DK, Endou H, et al. Novel liver-specific organic anion transporter OAT7 that operates the exchange of sulfate conjugates for short chain fatty acid butyrate. Hepatology. 2007;45(4):1046–55.

37. Mandal AK, Mount DB. The molecular physiology of uric acid homeostasis. Annu Rev Physiol. 2015;77:323–45.

38. Nakatochi M, Kanai M, Nakayama A, Hishida A, Kawamura Y, Ichihara S, et al. Genome-wide meta-analysis identifies multiple novel loci associated with serum uric acid levels in Japanese individuals. Commun Biol. 2019;2:115.

39. Eckardt KU, Alper SL, Antignac C, Bleyer AJ, Chauveau D, Dahan K, et al. Autosomal dominant tubulointerstitial kidney disease: diagnosis, classification, and management--A KDIGO consensus report. Kidney Int. 2015;88(4):676–83.

40. Kolz M, Johnson T, Sanna S, Teumer A, Vitart V, Perola M, et al. Meta-analysis of 28,141 individuals identifies common variants within five new loci that influence uric acid concentrations. PLoS Genet. 2009;5(6):e1000504.

41. Huang W, Shaikh SN, Ganapathy ME, Hopfer U, Leibach FH, Carter AL, et al. Carnitine transport and its inhibition by sulfonylureas in human kidney proximal tubular epithelial cells. Biochem Pharmacol. 1999;58(8):1361–70.

42. Bachhawat AK, Yadav S. The glutathione cycle: Glutathione metabolism beyond the gamma-glutamyl cycle. IUBMB Life. 2018;70(7):585–92.

43. Wang CK, Yang SC, Hsu SC, Chang FP, Lin YT, Chen SF, et al. CHAC2 is essential for self-renewal and glutathione maintenance in human embryonic stem cells. Free Radic Biol Med. 2017;113:439–51.

44. Frey IM, Rubio-Aliaga I, Siewert A, Sailer D, Drobyshev A, Beckers J, et al. Profiling at mRNA, protein, and metabolite levels reveals alterations in renal amino acid handling and glutathione metabolism in kidney tissue of Pept2-/- mice. Physiol Genomics. 2007;28(3):301–10.

45. Bahn A, Hagos Y, Reuter S, Balen D, Brzica H, Krick W, et al. Identification of a new urate and high affinity nicotinate transporter, hOAT10 (SLC22A13). J Biol Chem. 2008;283(24):16332–41.

46. Giri AK, Banerjee P, Chakraborty S, Kauser Y, Undru A, Roy S, et al. Genome wide association study of uric acid in Indian population and interaction of identified variants with Type 2 diabetes. Sci Rep. 2016;6:21440.

47. Houang EM, Sham YY, Bates FS, Metzger JM. Muscle membrane integrity in Duchenne muscular dystrophy: recent advances in copolymer-based muscle membrane stabilizers. Skelet Muscle. 2018;8(1):31.

48. Albrecht DE, Sherman DL, Brophy PJ, Froehner SC. The ABCA1 cholesterol transporter associates with one of two distinct dystrophin-based scaffolds in Schwann cells. Glia. 2008;56(6):611–8.

49. Haenggi T, Schaub MC, Fritschy JM. Molecular heterogeneity of the dystrophin-associated protein complex in the mouse kidney nephron: differential alterations in the absence of utrophin and dystrophin. Cell Tissue Res. 2005;319(2):299–313.

50. Mandal AK, Mercado A, Foster A, Zandi-Nejad K, Mount DB. Uricosuric targets of tranilast. Pharmacol Res Perspect. 2017;5(2):e00291.

51. Riedmaier AE, Burk O, van Eijck BAC, Schaeffeler E, Klein K, Fehr S, et al. Variability in hepatic expression of organic anion transporter 7/SLC22A9, a novel pravastatin uptake transporter: impact of genetic and regulatory factors. Pharmacogenomics J. 2016;16(4):341–51.

52. Mattila J, Havula E, Suominen E, Teesalu M, Surakka I, Hynynen R, et al. Mondo-Mlx Mediates Organismal Sugar Sensing through the Gli-Similar Transcription Factor Sugarbabe. Cell reports. 2015;13(2):350–64.

53. Ortega-Prieto P, Postic C. Carbohydrate Sensing Through the Transcription Factor ChREBP. Front Genet. 2019;10:472.

54. Stoltzman CA, Peterson CW, Breen KT, Muoio DM, Billin AN, Ayer DE. Glucose sensing by MondoA:Mlx complexes: a role for hexokinases and direct regulation of thioredoxin-interacting protein expression. Proc Natl Acad Sci U S A. 2008;105(19):6912–7.

55. Gosling AL, Boocock J, Dalbeth N, Harre Hindmarsh J, Stamp LK, Stahl EA, et al. Mitochondrial genetic variation and gout in Maori and Pacific people living in Aotearoa New Zealand. Ann Rheum Dis. 2018;77(4):571–8.

56. Menezes MJ, Guo Y, Zhang J, Riley LG, Cooper ST, Thorburn DR, et al. Mutation in mitochondrial ribosomal protein S7 (MRPS7) causes congenital sensorineural deafness, progressive hepatic and renal failure and lactic acidemia. Hum Mol Genet. 2015;24(8):2297–307.

57. Dogra R, Bhatia R, Shankar R, Bansal P, Rawal RK. Enasidenib: First Mutant IDH2 Inhibitor for the Treatment of Refractory and Relapsed Acute Myeloid Leukemia. Anticancer Agents Med Chem. 2018;18(14):1936–51.

58. Amary MF, Damato S, Halai D, Eskandarpour M, Berisha F, Bonar F, et al. Ollier disease and Maffucci syndrome are caused by somatic mosaic mutations of IDH1 and IDH2. Nat Genet. 2011;43(12):1262–5.

59. Consortium GP. A global reference for human genetic variation. Nature. 2015;526(7571):68–74.

60. Yang J, Lee SH, Goddard ME, Visscher PM. GCTA: a tool for genome-wide complex trait analysis. Am J Hum Genet. 2011;88(1):76–82.

61. Bulik-Sullivan BK, Loh PR, Finucane HK, Ripke S, Yang J, Schizophrenia Working Group of the Psychiatric Genomics C, et al. LD Score regression distinguishes confounding from polygenicity in genome-wide association studies. Nat Genet. 2015;47(3):291–5.

62. Yang J, Ferreira T, Morris AP, Medland SE, Genetic Investigation of ATC, Replication DIG, et al. Conditional and joint multiple-SNP analysis of GWAS summary statistics identifies additional variants influencing complex traits. Nat Genet. 2012;44(4):369–75, S1-3.

63. Gao XY, Stamier J, Martin ER. A multiple testing correction method for genetic association studies using correlated single nucleotide polymorphisms. Genet epidemiol. 2008;32(4):361–9.

64. Finucane HK, Bulik-Sullivan B, Gusev A, Trynka G, Reshef Y, Loh PR, et al. Partitioning heritability by functional annotation using genome-wide association summary statistics. Nat Genet. 2015;47(11):1228.

65. Benjamini Y, Hochberg Y. Controlling the False Discovery Rate - a Practical and Powerful Approach to Multiple Testing. J R Stat Soc B. 1995;57(1):289–300.

66. Wickham H. ggplot2: elegant graphics for data analysis: Springer; 2016.

67. Kichaev G, Roytman M, Johnson R, Eskin E, Lindstrom S, Kraft P, et al. Improved methods for multi-trait fine mapping of pleiotropic risk loci. Bioinformatics. 2017;33(2):248–55.

68. Giambartolomei C, Vukcevic D, Schadt EE, Franke L, Hingorani AD, Wallace C, et al. Bayesian Test for Colocalisation between Pairs of Genetic Association Studies Using Summary Statistics. Plos Genetics. 2014;10(5):e1004383.

69. Fadason T, Ekblad C, Ingram JR, Schierding WS, O’Sullivan JM. Physical Interactions and Expression Quantitative Traits Loci Identify Regulatory Connections for Obesity and Type 2 Diabetes Associated SNPs. Front Genet. 2017;8:150.

70. Rao SS, Huntley MH, Durand NC, Stamenova EK, Bochkov ID, Robinson JT, et al. A 3D map of the human genome at kilobase resolution reveals principles of chromatin looping. Cell. 2014;159(7):1665–80.

71. GTEx Consortium. The Genotype-Tissue Expression (GTEx) project. Nat Genet. 2013;45(6):580–5.

72. Cadzow M, Merriman TR, Dalbeth N. Performance of gout definitions for genetic epidemiological studies: analysis of UK Biobank. Arthritis Res Ther. 2017;19(1):181.

73. Liman ER, Tytgat J, Hess P. Subunit stoichiometry of a mammalian K+ channel determined by construction of multimeric cDNAs. Neuron. 1992;9(5):861–71.

