## Supplementary Figures for "Genomic dissection of 43 serum urate-associated loci provides multiple insights into molecular mechanisms of urate control"

### Figure S1

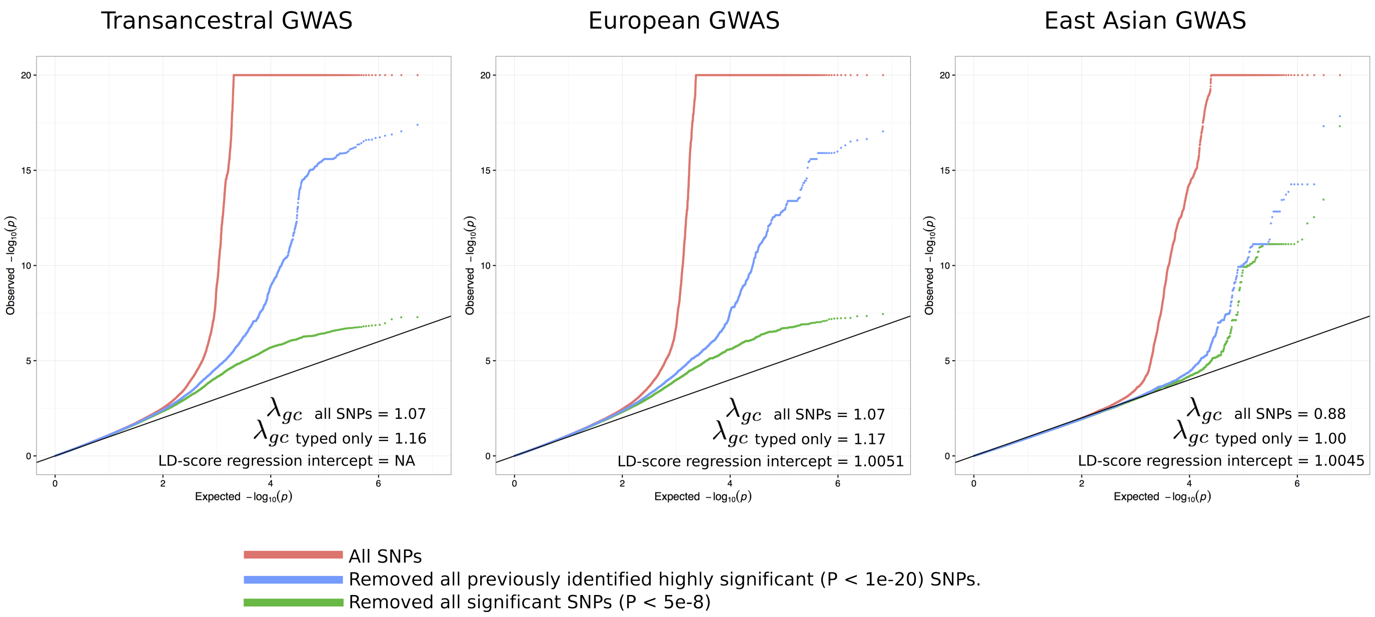
Transancestral GWAS
European GWAS
East Asian GWAS
Observed −log10(p)
Expected −log10(p)
0
2
4
6
5
10
15
20
λgc all SNPs = 1.07
λgc typed only = 1.16
LD-score regression intercept = NA
λgc all SNPs = 1.07
λgc typed only = 1.17
LD-score regression intercept = 1.0051
λgc all SNPs = 0.88
λgc typed only = 1.00
LD-score regression intercept = 1.0045
All SNPs
Removed all previously identified highly significant (P < 1e-20) SNPs.
Removed all significant SNPs (P < 5e-8)

### Figure S2

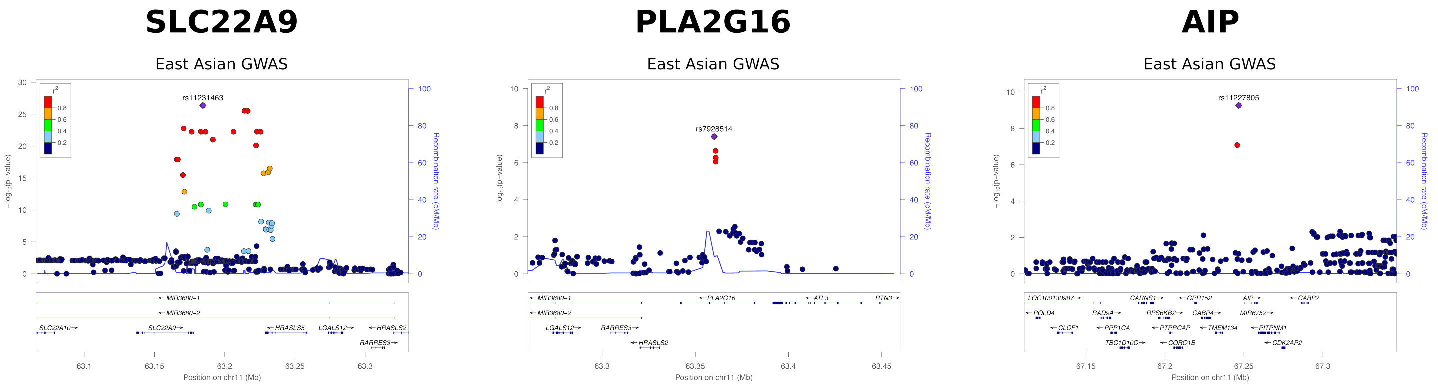
SLC22A9
East Asian GWAS
rs11231463
r²
0.8
0.6
0.4
0.2
−log₁₀(p−value)
Recombination rate (cM/Mb)
←MIR3680−1
←MIR3680−2
SLC22A10→
SLC22A9→
←HRASLS5
LGALS12→
←HRASLS2
RARRES3→
63.1
63.15
63.2
63.25
63.3
Position on chr11 (Mb)
PLA2G16
East Asian GWAS
rs7928514
←MIR3680−1
←MIR3680−2
LGALS12→
RARRES3→
←HRASLS2
←PLA2G16
←ATL3
RTN3→
63.3
63.35
63.4
63.45
Position on chr11 (Mb)
AIP
East Asian GWAS
rs11227805
LOC100130987→
CARNS1→
←GPR152
AIP→
←CABP2
←POLD4
RAD9A→
RPS6KB2→
CABP4→
MIR6752→
←CLCF1
←PPP1CA
←PTPRCAP
←TMEM134
←PITPNM1
TBC1D10C→
←CORO1B
←CDK2AP2
67.15
67.2
67.25
67.3
Position on chr11 (Mb)

### Figure S3

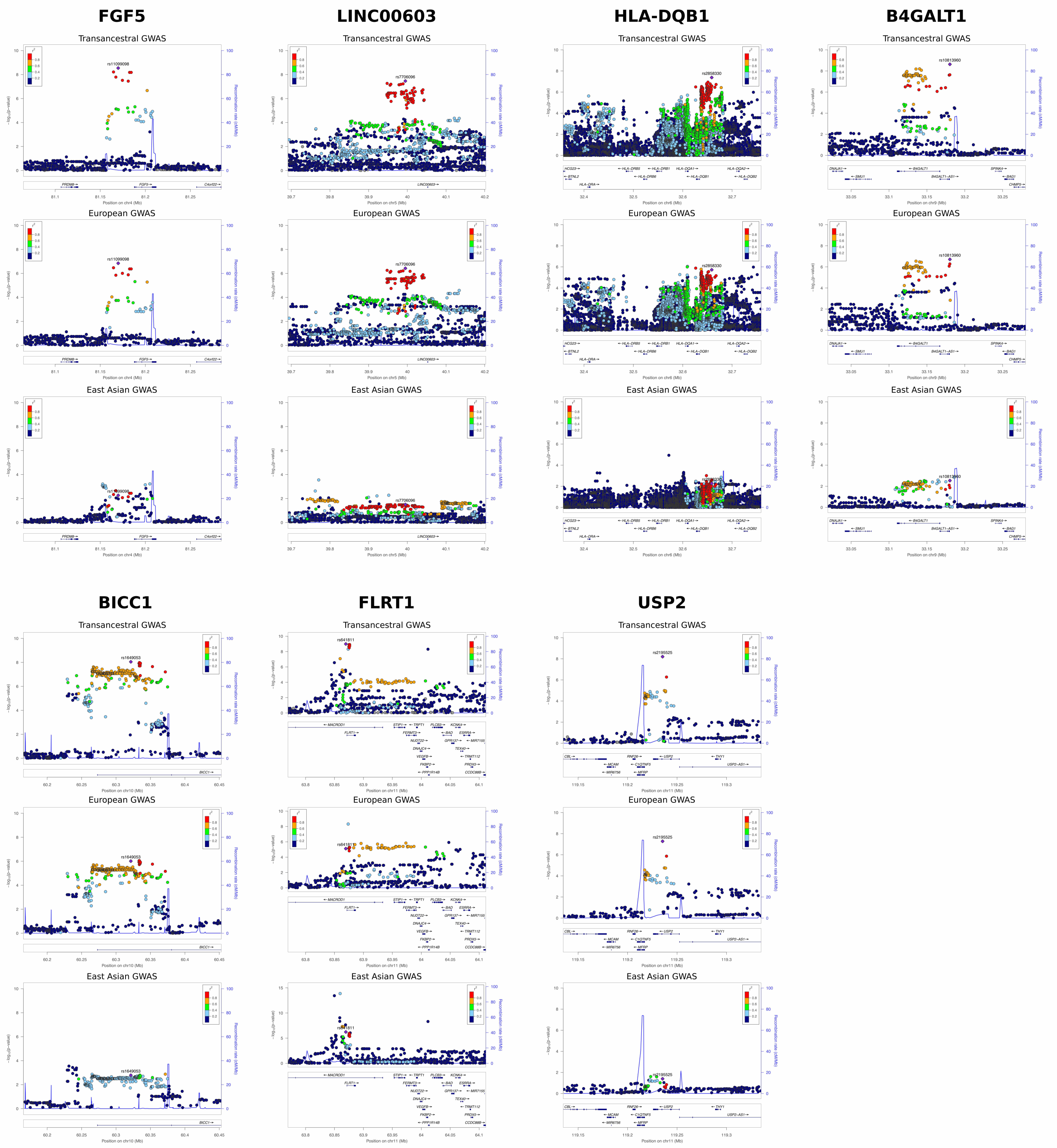
FGF5
LINC00603
HLA-DQB1
B4GALT1
Transancestral GWAS
European GWAS
East Asian GWAS
BICC1
FLRT1
USP2
Transancestral GWAS
European GWAS
East Asian GWAS

### Figure S4

a) **rs12289836** (MAF EUR=0.35, MAF EAS=0.22)

European GWAS | East Asian GWAS

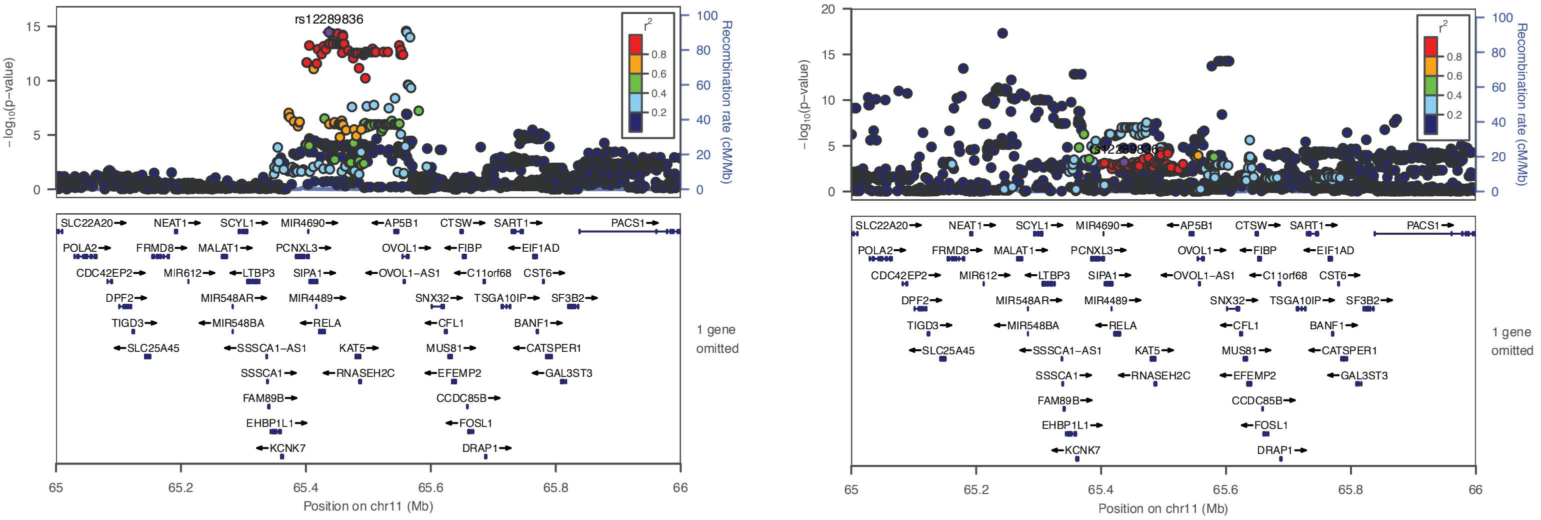


b) **rs11227200** (MAF EUR=0.26, MAF EAS=0.14)

European GWAS | East Asian GWAS

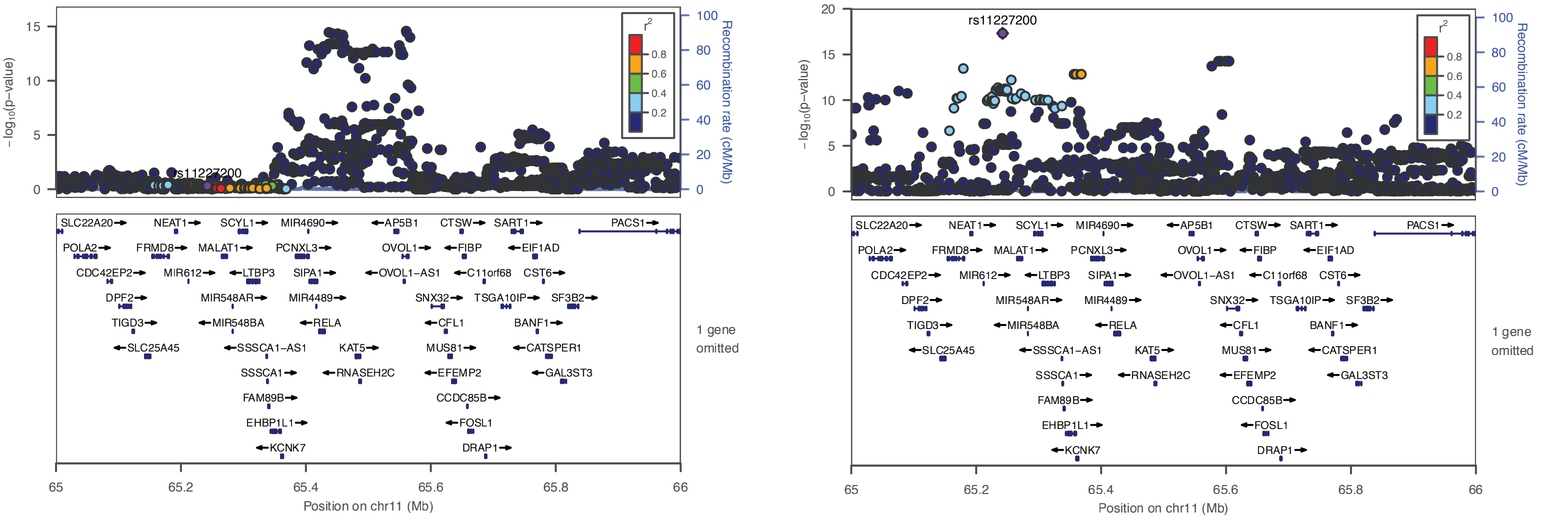

### Figure S5

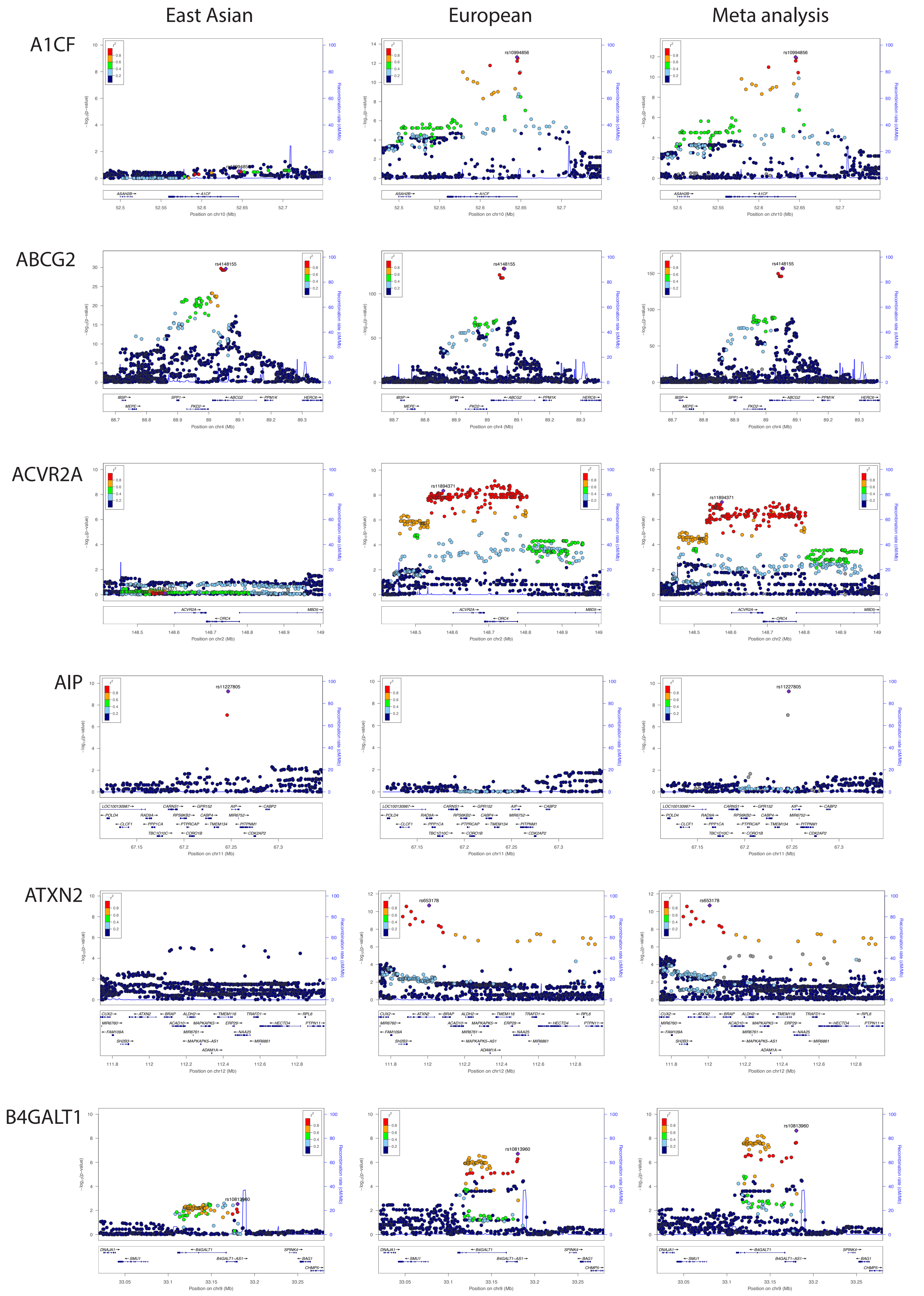

East Asian
European
Meta analysis
A1CF
ABCG2
ACVR2A
AIP
ATXN2
B4GALT1
rs10994856
rs4148155
rs11894371
rs11227805
rs653178
rs10813960
−log10(p-value)
Recombination rate (cM/Mb)
Position on chr10 (Mb)
Position on chr4 (Mb)
Position on chr2 (Mb)
Position on chr11 (Mb)
Position on chr12 (Mb)
Position on chr9 (Mb)

### Figure S5 cont.

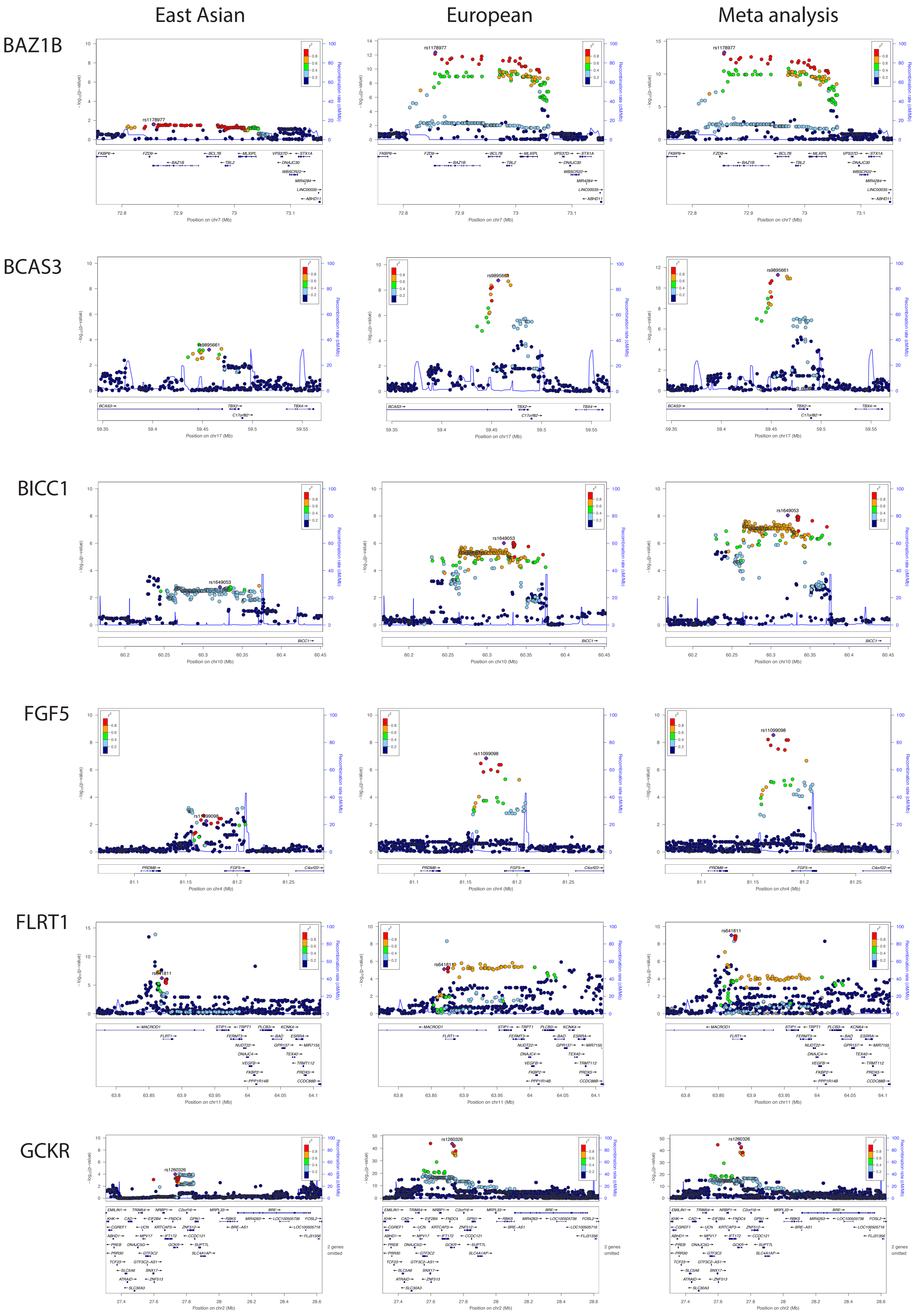

### Figure S5 cont.

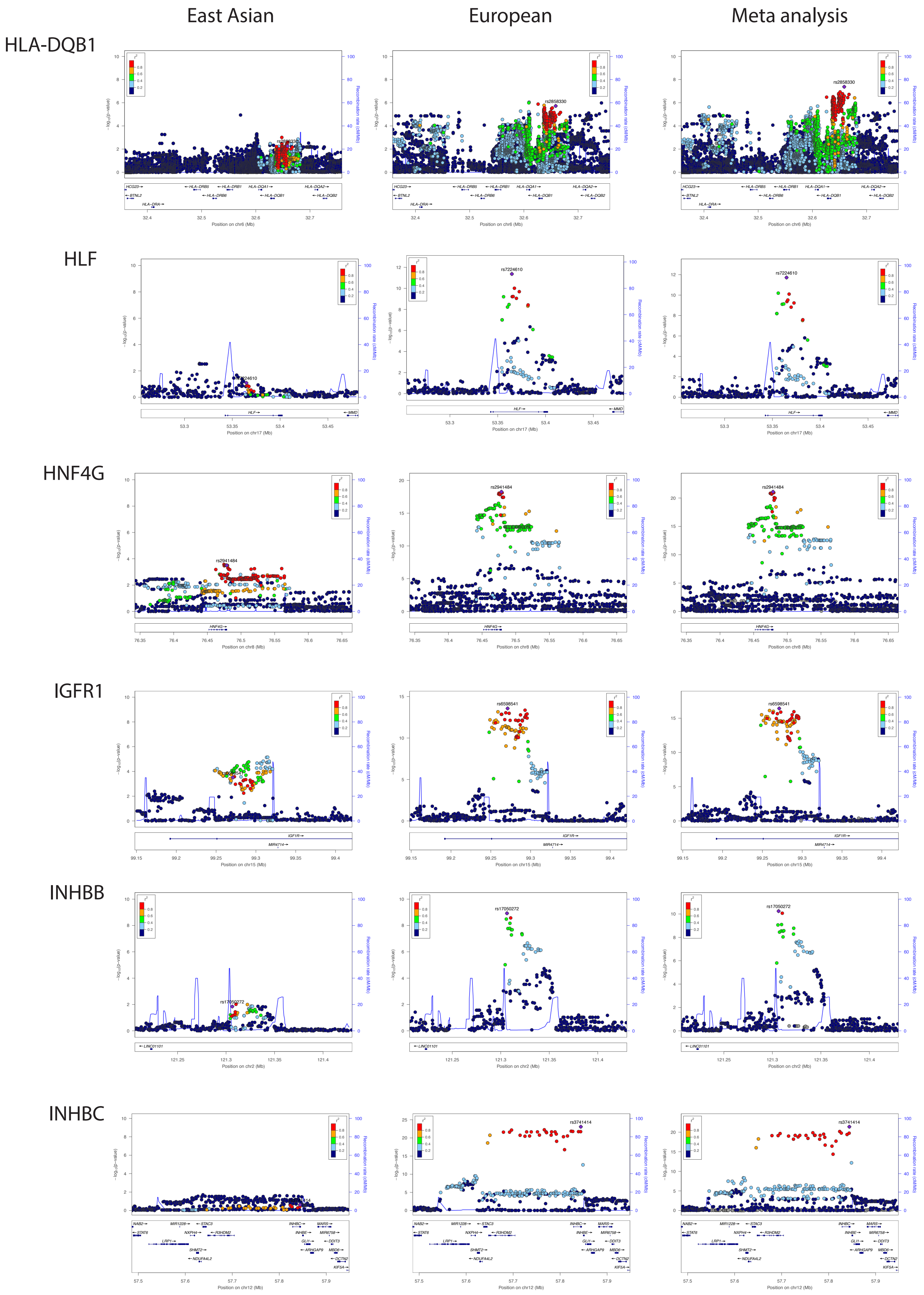

Figure S5 cont.

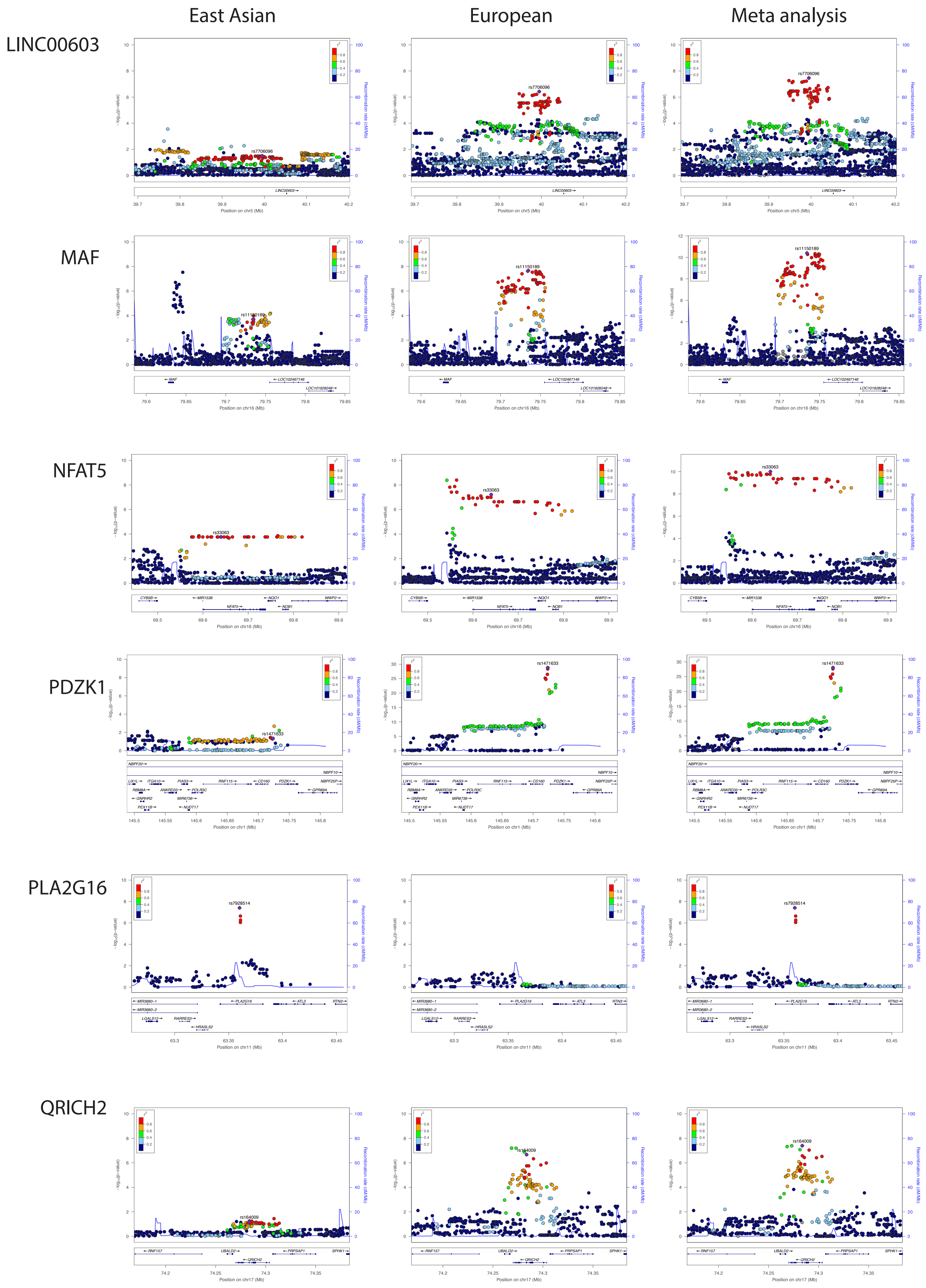

### Figure S5 cont.

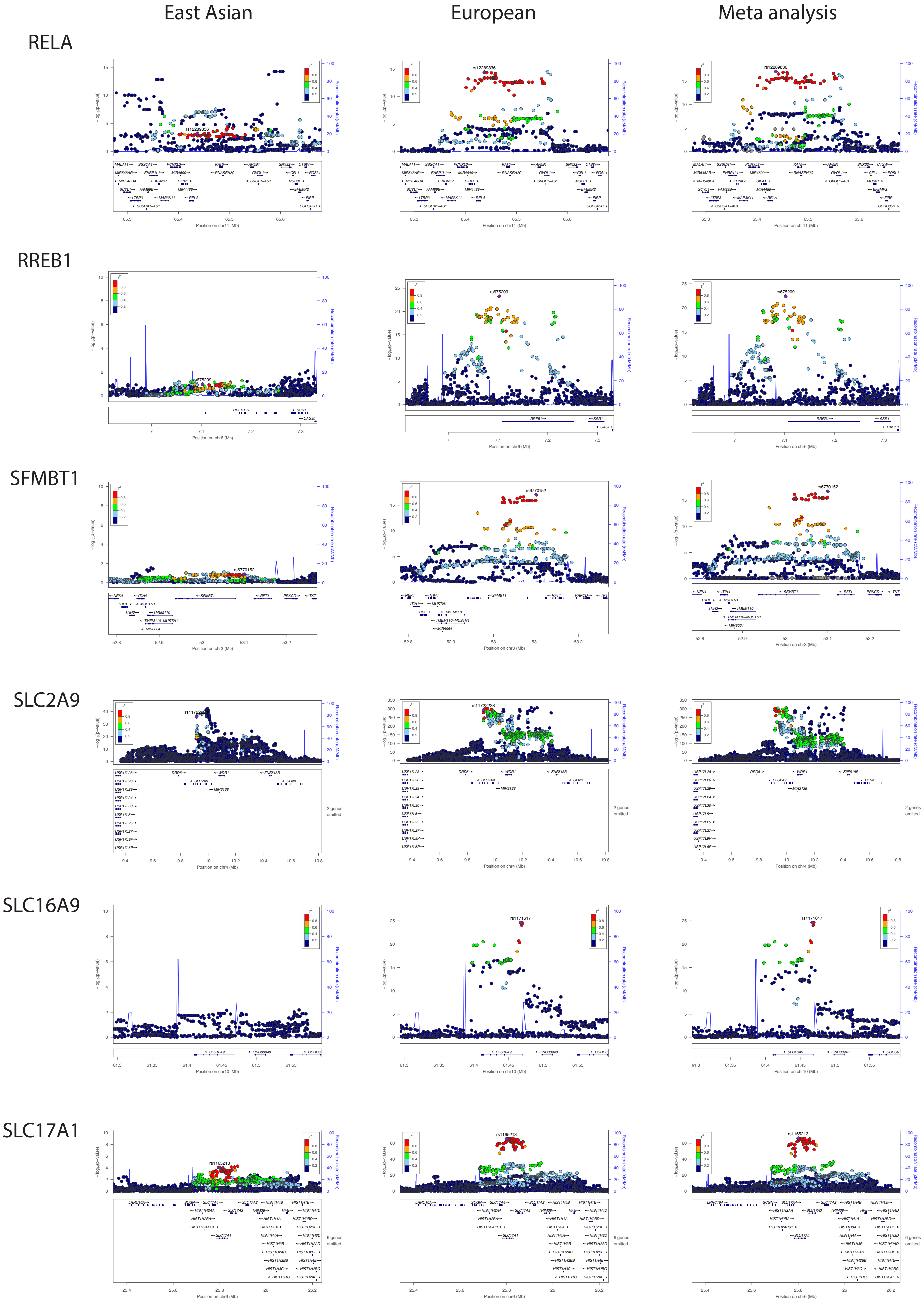

Figure S5 cont.

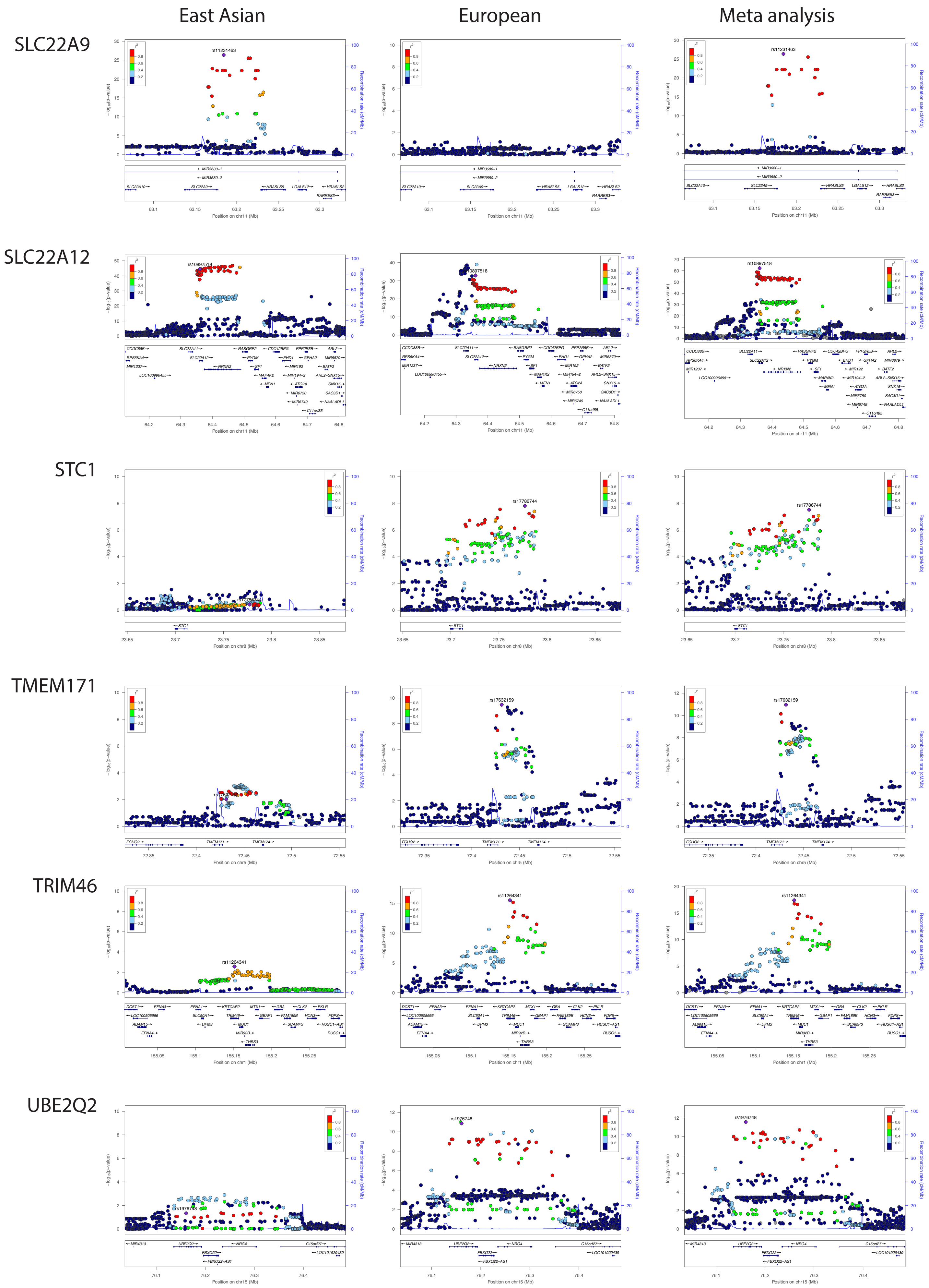

Figure S5 cont.

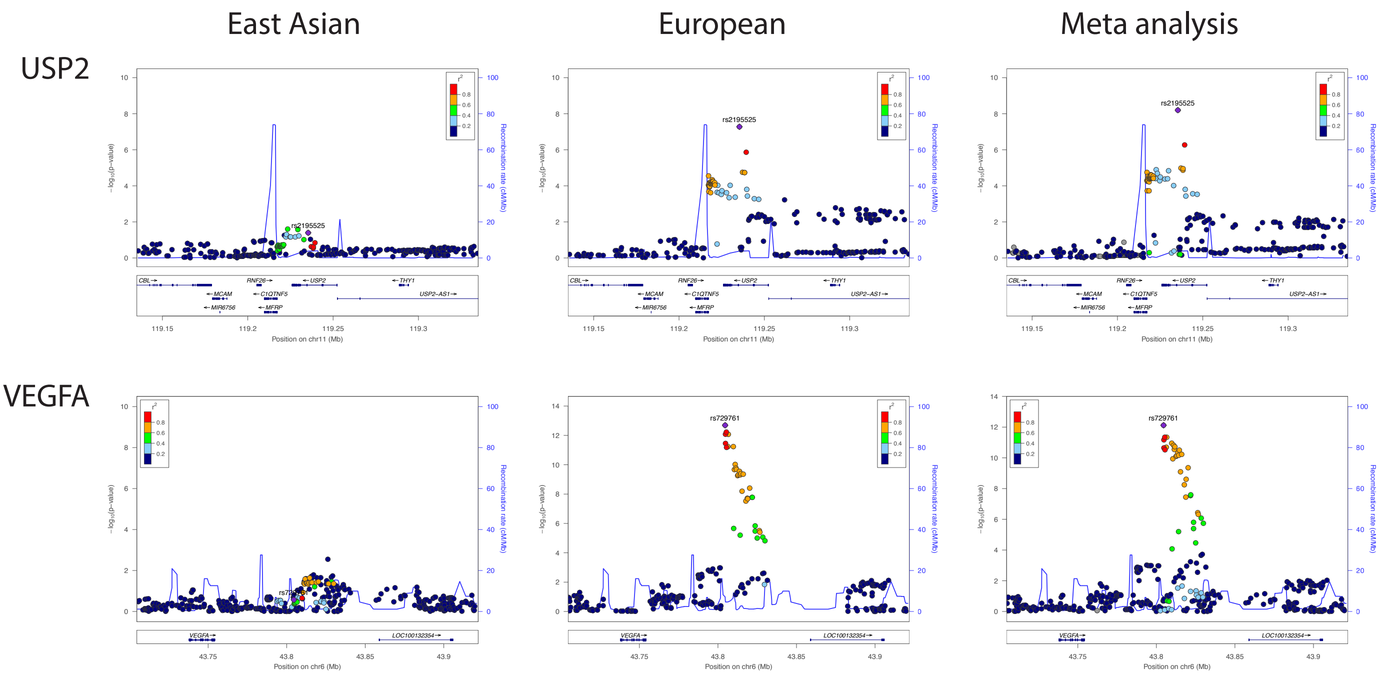

Figure S6

a) **rs1150189** (MAF EUR=0.30, MAF EAS=0.18)

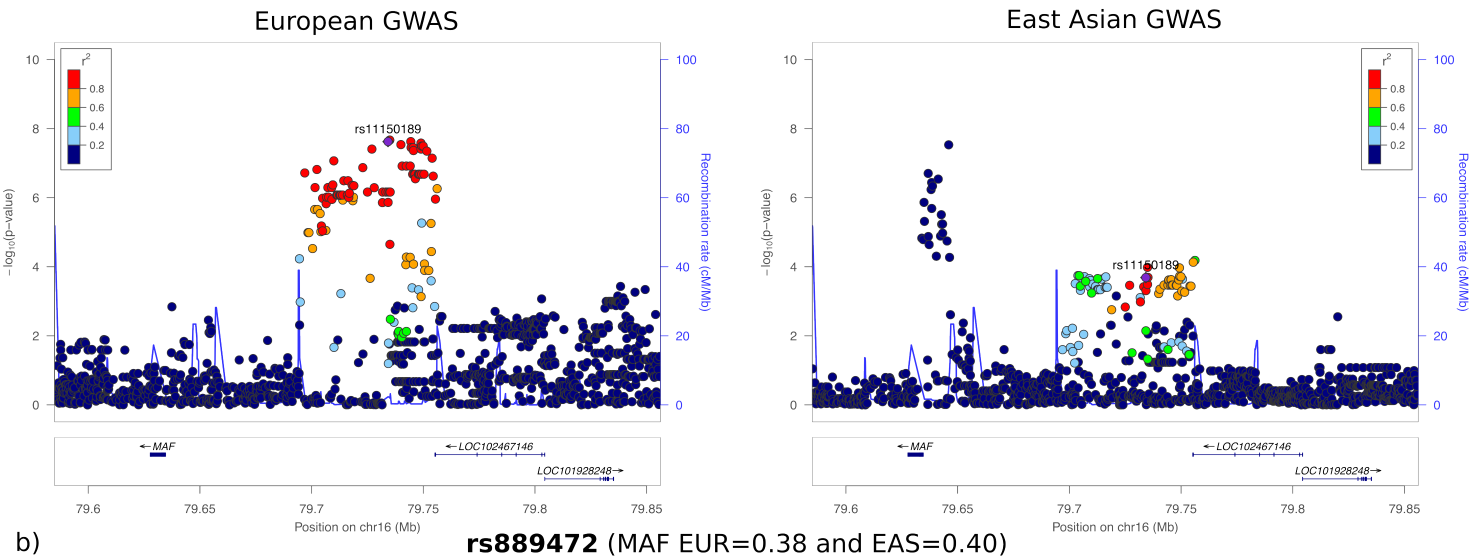


b) **rs889472** (MAF EUR=0.38 and EAS=0.40)

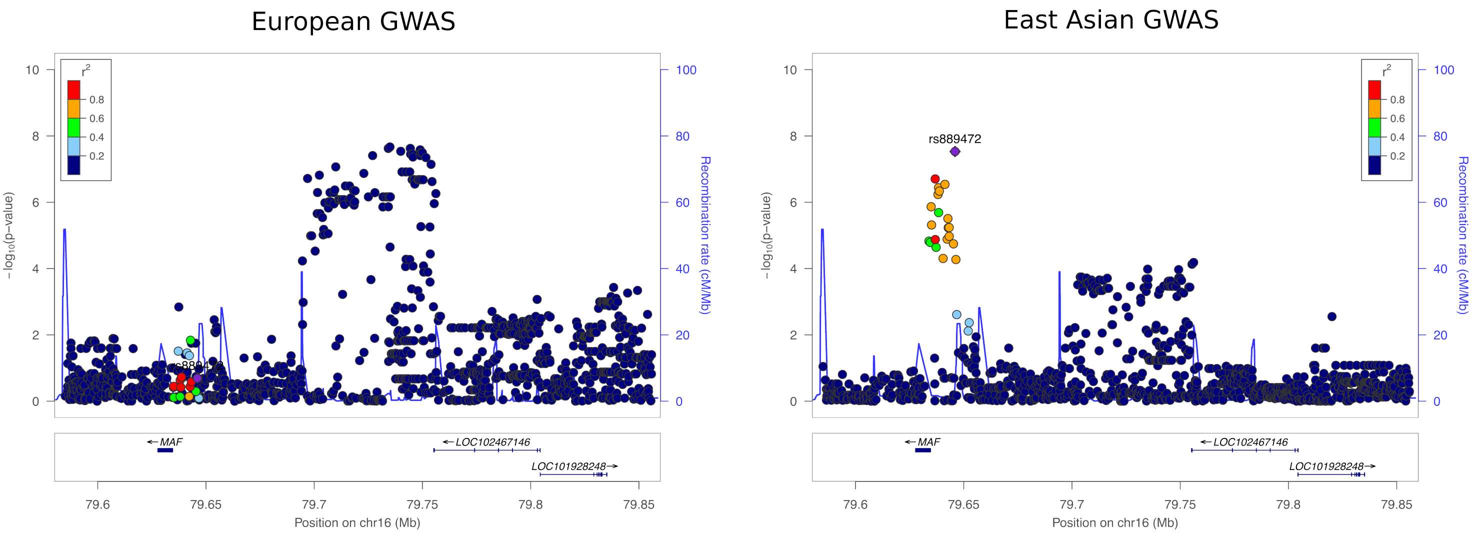

### Figure S7

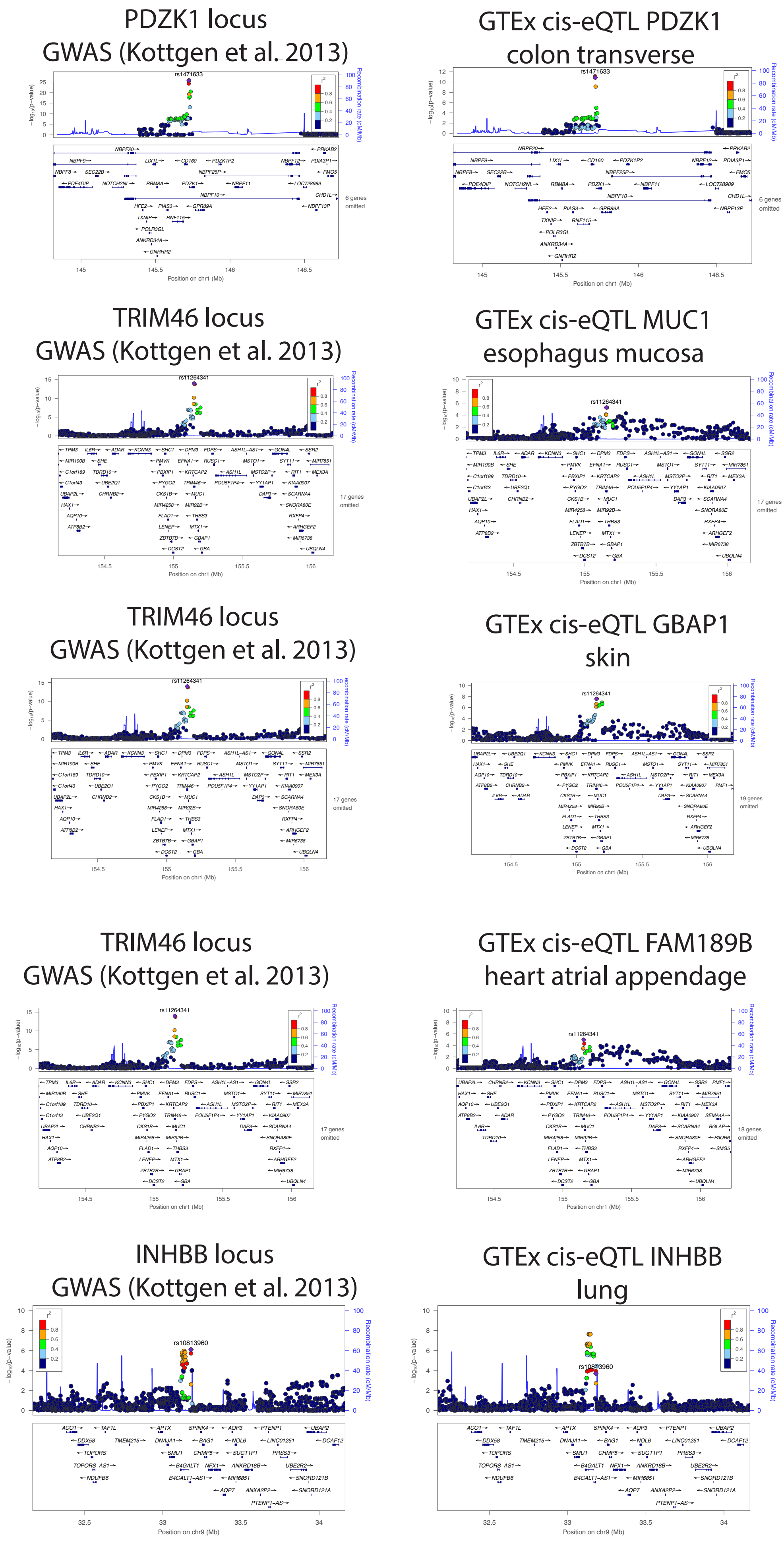
PDZK1 locus
GWAS (Kottgen et al. 2013)
GTEx cis-eQTL PDZK1
colon transverse
rs1471633
TRIM46 locus
GWAS (Kottgen et al. 2013)
GTEx cis-eQTL MUC1
esophagus mucosa
rs11264341
TRIM46 locus
GWAS (Kottgen et al. 2013)
GTEx cis-eQTL GBAP1
skin
TRIM46 locus
GWAS (Kottgen et al. 2013)
GTEx cis-eQTL FAM189B
heart atrial appendage
INHBB locus
GWAS (Kottgen et al. 2013)
GTEx cis-eQTL INHBB
lung
rs10813960
Position on chr1 (Mb)
Position on chr9 (Mb)
Recombination rate (cM/Mb)
−log10(p-value)

### Figure S7 cont.

#### SFMBT1 locus
#### GWAS (Kottgen et al. 2013)

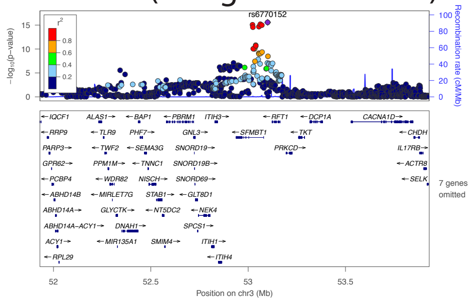


#### GTEx cis-eQTL TMEM110
#### adipose subcutaneous

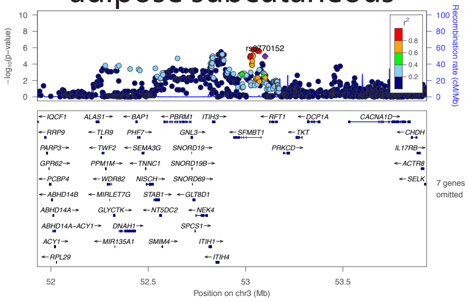


#### SFMBT1 locus
#### GWAS (Kottgen et al. 2013)

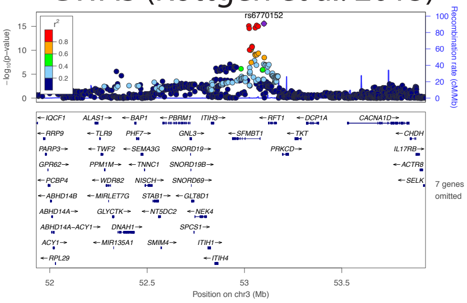


#### GTEx cis-eQTL SFMBT1
#### colon transverse

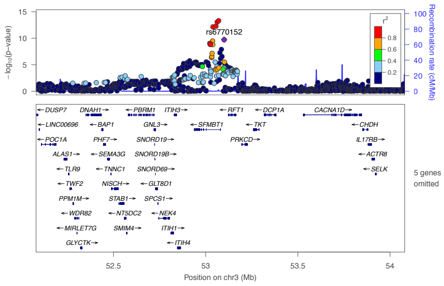


#### HLA-DQB1 locus
#### GWAS (Kottgen et al. 2013)

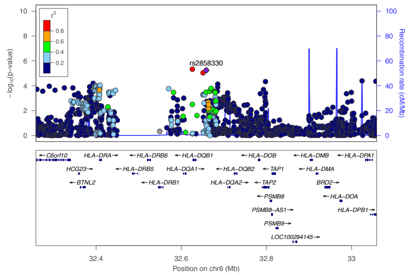


#### GTEx cis-eQTL HLA-DQA2
#### prostate

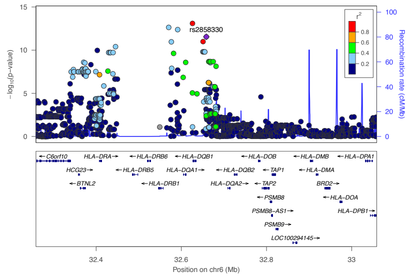


#### BAZ1B locus
#### GWAS (Kottgen et al. 2013)

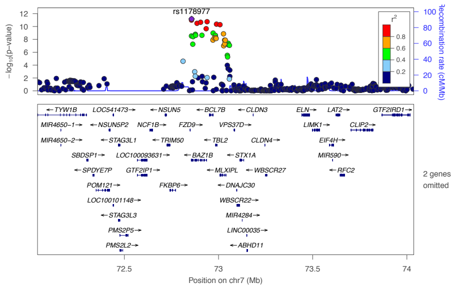


#### GTEx cis-eQTL MLXIPL
#### transformed fibroblasts

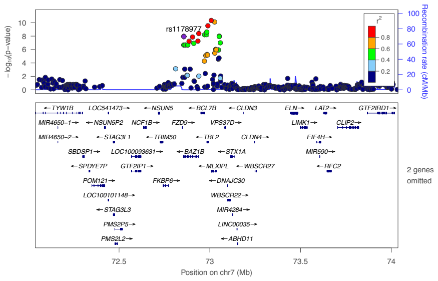


#### B4GALT1 locus
#### GWAS (Kottgen et al. 2013)

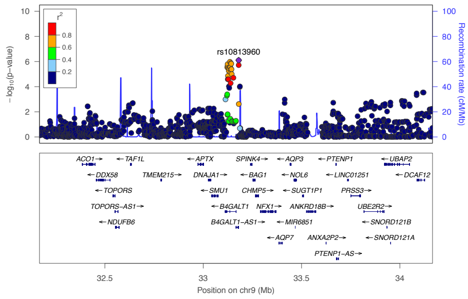


#### GTEx cis-eQTL B4GALT1
#### EBV transformed lymphocytes

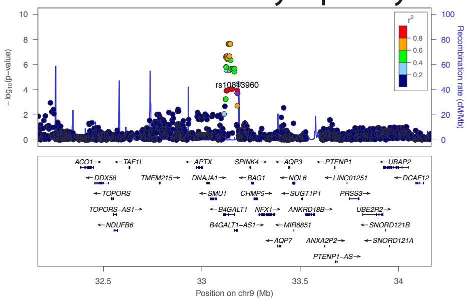

Figure S7 cont.

SLC16A9 locus
GWAS (Kottgen et al. 2013)

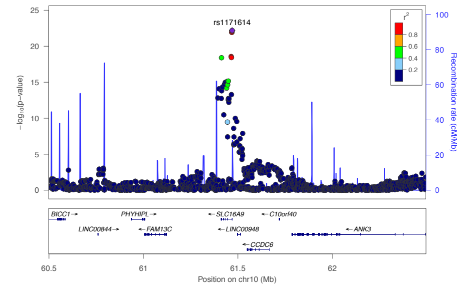


GTEx cis-eQTL SLC16A9
thyroid

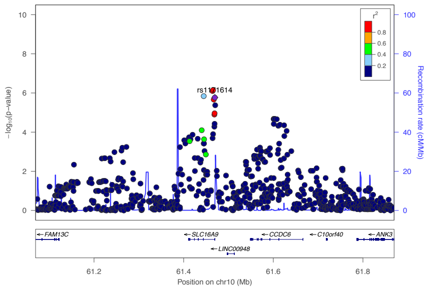


RELA locus
GWAS (Kottgen et al. 2013)

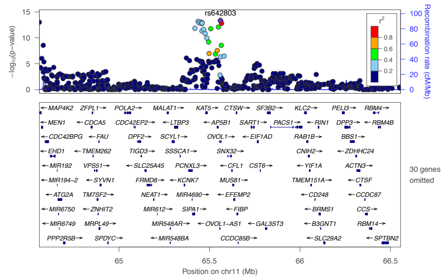


GTEx cis-eQTL OVOL1-AS1
nucleus accumbens basal ganglia

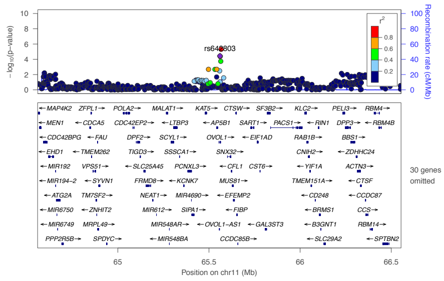


INHBC locus
GWAS (Kottgen et al. 2013)

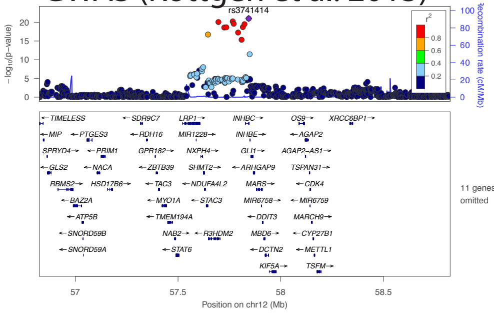


GTEx cis-eQTL R3HDM2
transformed fibroblasts

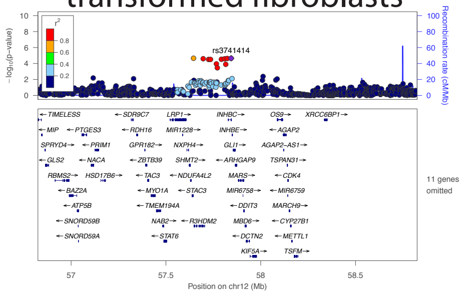


UBE2Q2 locus
GWAS (Kottgen et al. 2013)

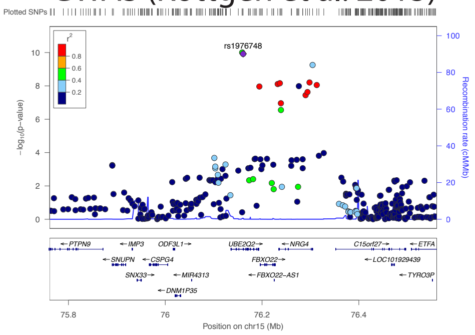


GTEx cis-eQTL UBE2Q2
cortex

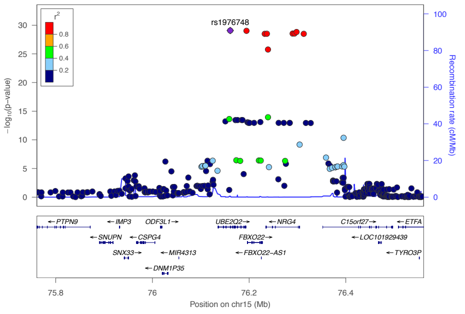


IGF1R locus
GWAS (Kottgen et al. 2013)

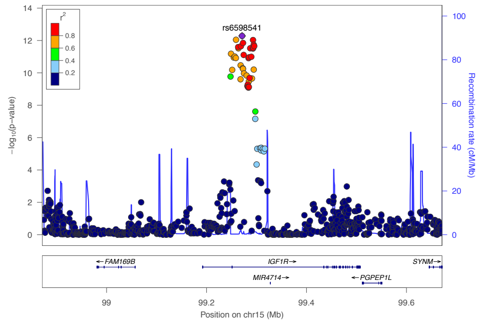


GTEx cis-eQTL IGF1R
heart left ventricle

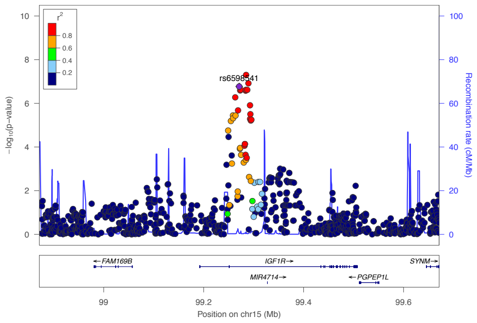

### Figure S7 cont.

#### MAF locus
#### GWAS (Kottgen et al. 2013)

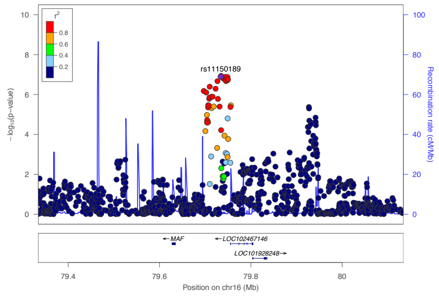


#### GTEx cis-eQTL MAFTRR
#### colon sigmoid

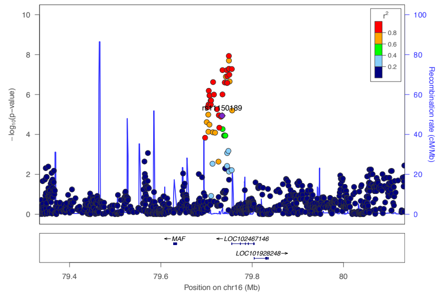


#### QRICH2 locus
#### GWAS (Kottgen et al. 2013)

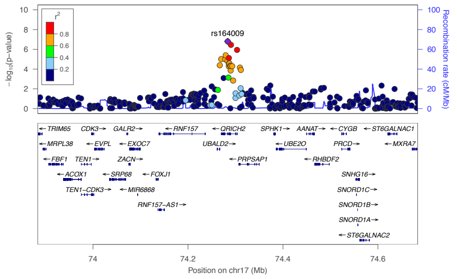


#### GTEx trans-eQTL UBALD2
#### esophagus muscularis

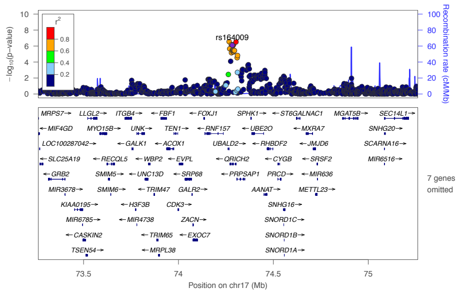


#### QRICH2 locus
#### GWAS (Kottgen et al. 2013)

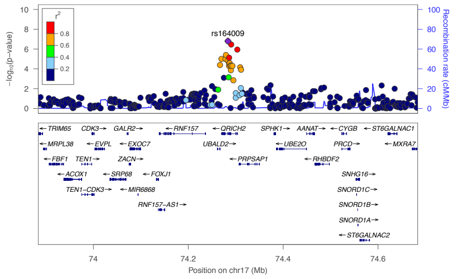


#### GTEx cis-eQTL PRPSAP1
#### anterior cingulate cortex

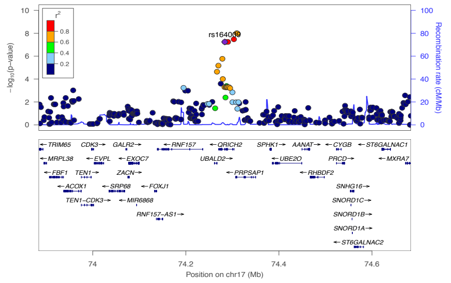


#### DHRS9 locus
#### GWAS (Kottgen et al. 2013)

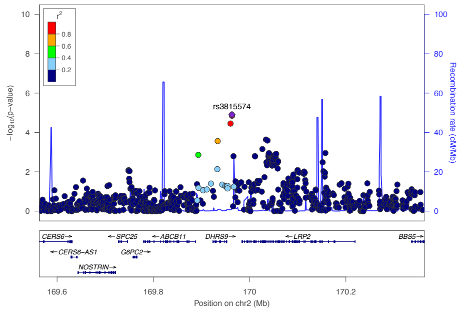


#### GTEx cis-eQTL DHRS9
#### whole blood

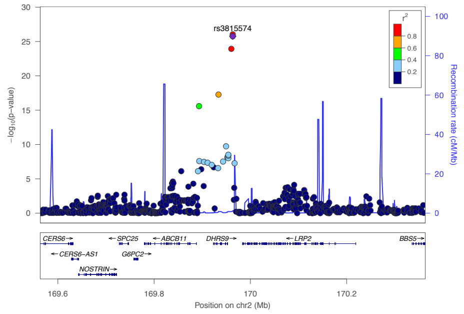


#### RAI14 locus
#### GWAS (Kottgen et al. 2013)

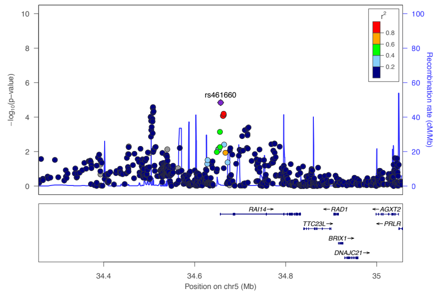


#### GTEx cis-eQTL RAI14
#### thyroid

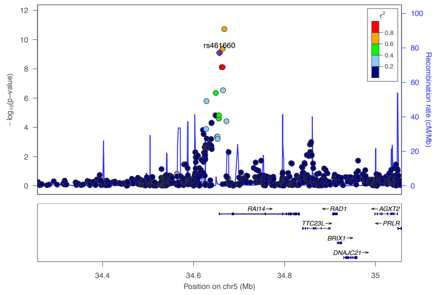

### Figure S7 cont.

#### MLXIP locus
#### GWAS (Kottgen et al. 2013)

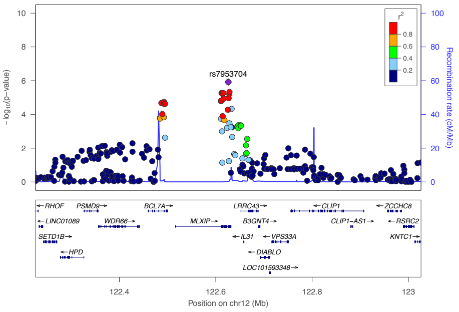


#### GTEx cis-eQTL MLXIP
#### small intestine

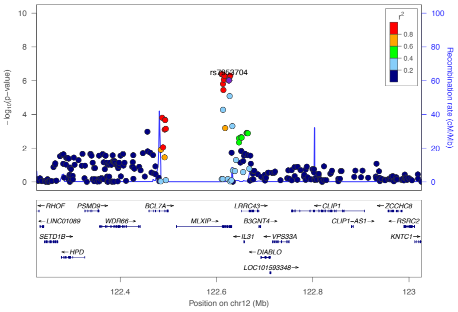


#### IDH2 locus
#### GWAS (Kottgen et al. 2013)

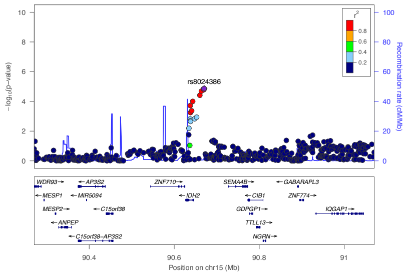


#### GTEx cis-eQTL IDH2
#### heart atrial appendage

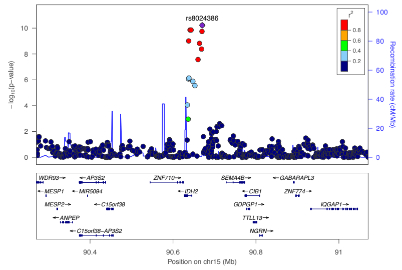


#### MRPS7 locus
#### GWAS (Kottgen et al. 2013)

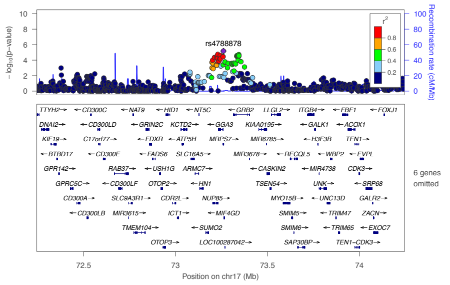


#### GTEx cis-eQTL GGA3
#### thyroid

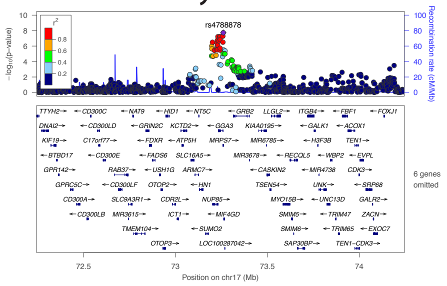


#### MRPS7 locus
#### GWAS (Kottgen et al. 2013)

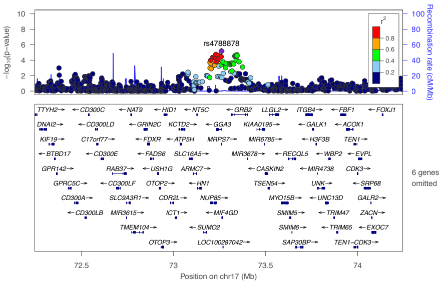


#### GTEx cis-eQTL MRPS7
#### dorsolateral cortex

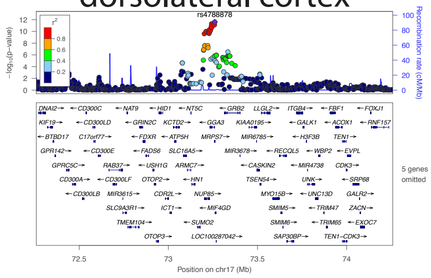


#### NFAT5 locus
#### GWAS (Kottgen et al. 2013)

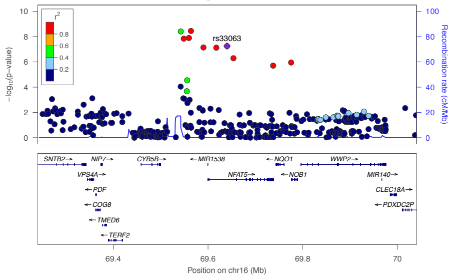


#### GTEx trans-eQTL A1FL
#### brain substantial nigra

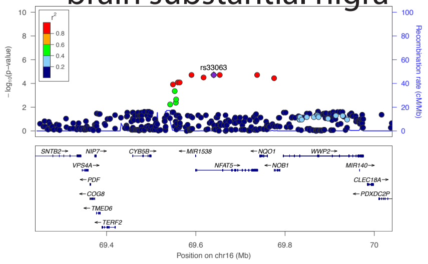

Figure S7 cont.

NFAT5 locus
GWAS (Kottgen et al. 2013)

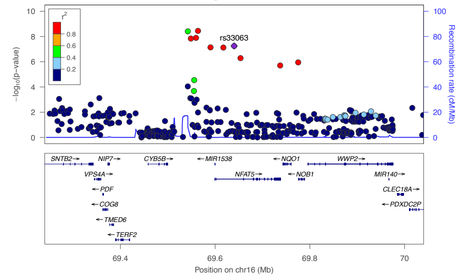


GTEx trans-eQTL CACNA2D3
basal ganglia

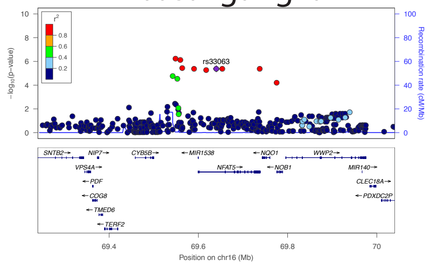


NFAT5 locus
GWAS (Kottgen et al. 2013)

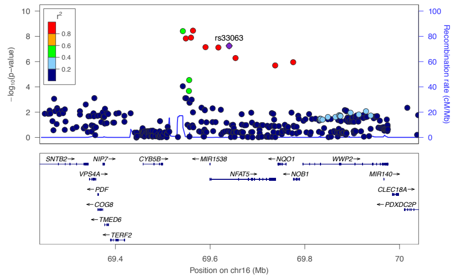


GTEx trans-eQTL STIM1
basal ganglia

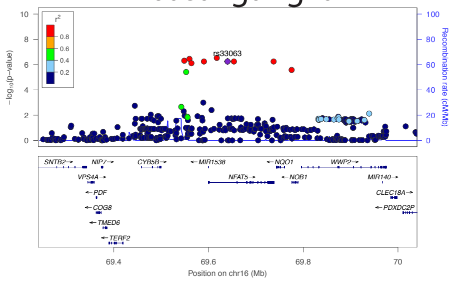


SLC16A9 locus
GWAS (Kottgen et al. 2013)

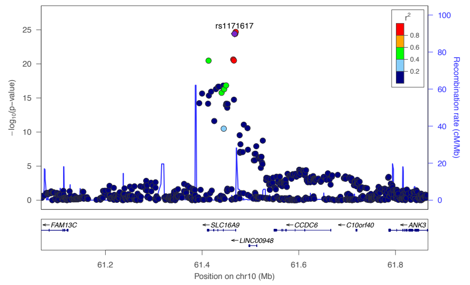


GTEx trans-eQTL ANKS1B
testis

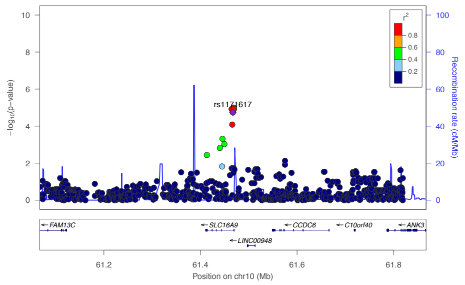


SLC16A9 locus
GWAS (Kottgen et al. 2013)

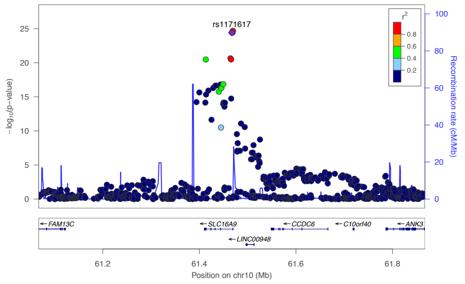


GTEx trans-eQTL DSCAM
brain hypothalamus

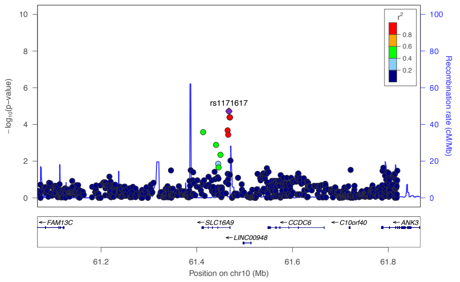


BAZ1B locus
GWAS (Kottgen et al. 2013)

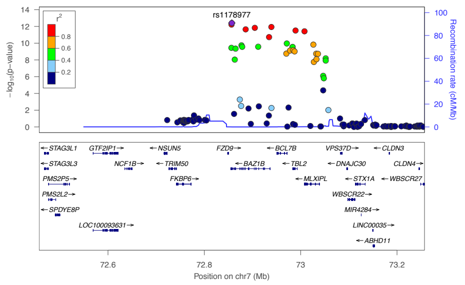


GTEx trans-eQTL RNF24
brain cortex

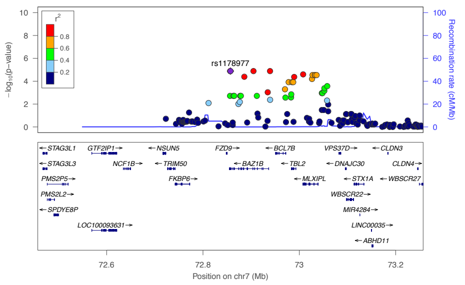

### Figure S7 cont.

#### QRICH2 locus
#### GWAS (Kottgen et al. 2013)

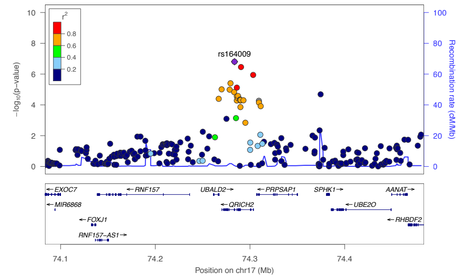


#### GTEx trans-eQTL PPP3R1
#### heart left ventricle

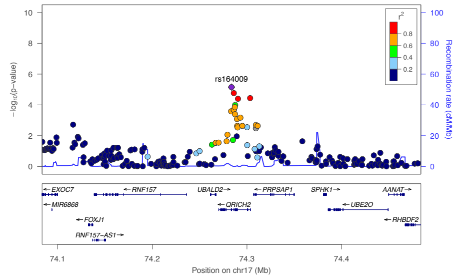


#### INHBB locus
#### GWAS (Kottgen et al. 2013)

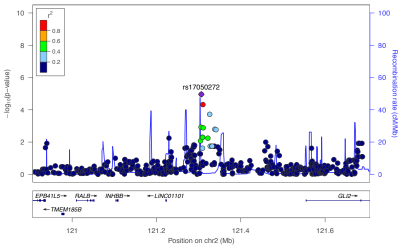


#### GTEx trans-eQTL CHAC2
#### brain caudate basal ganglia

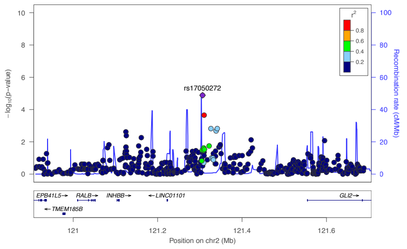


#### INHBB locus
#### GWAS (Kottgen et al. 2013)

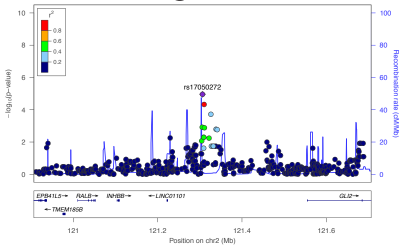


#### GTEx trans-eQTL ZNF804A
#### brain anterior cingulate cortex

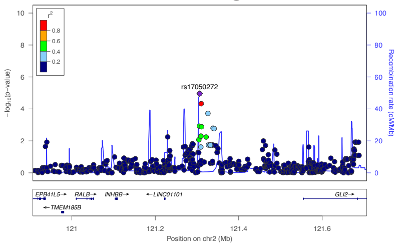


#### UBE2Q2 locus
#### GWAS (Kottgen et al. 2013)

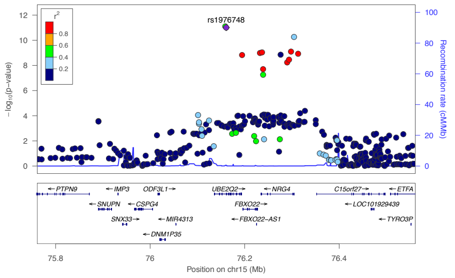


#### GTEx trans-eQTL COL11A1
#### colon transverse

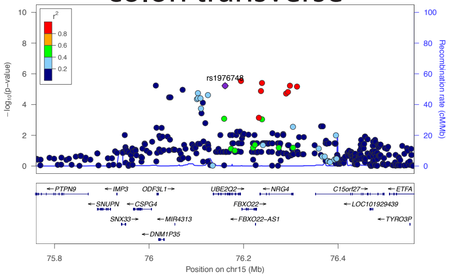


#### HNF4G locus
#### GWAS (Kottgen et al. 2013)

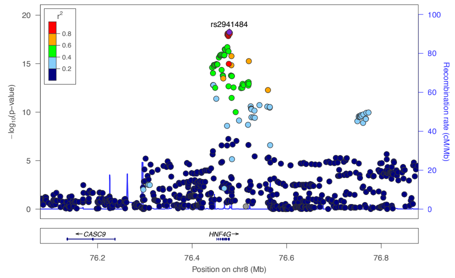


#### GTEx trans-eQTL CSMD2
#### brain cerebellum

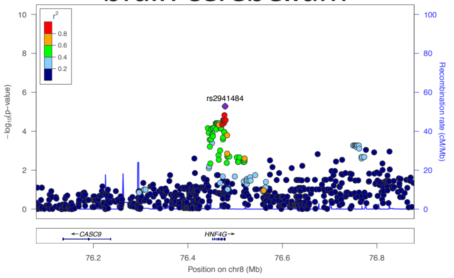

### Figure S7 cont.

#### INHBC locus
#### GWAS (Kottgen et al. 2013)

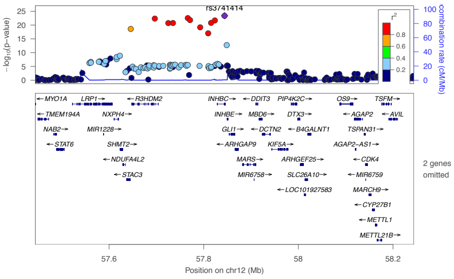


#### GTEx trans-eQTL SPIN1
#### brain frontal cortex ba9

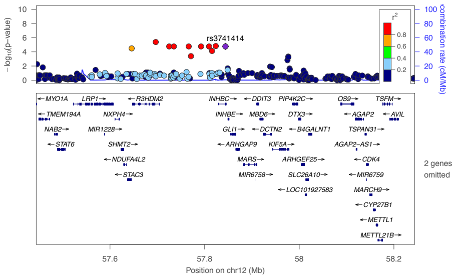


#### DHRS9 locus
#### GWAS (Kottgen et al. 2013)

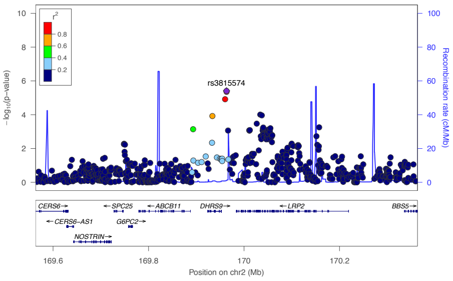


#### GTEx trans-eQTL JHDM1D
#### brain hippocampus

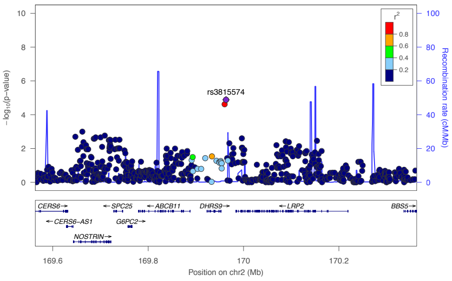


#### RREB1 locus
#### GWAS (Kottgen et al. 2013)

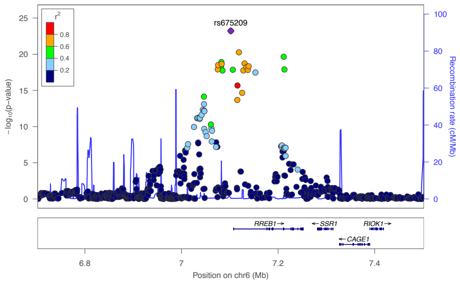


#### GTEx trans-eQTL UTRN
#### brain putamen basal ganglia

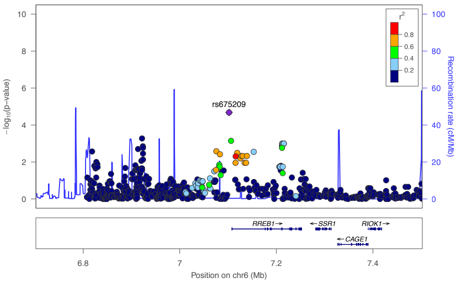


#### HLF locus
#### GWAS (Kottgen et al. 2013)

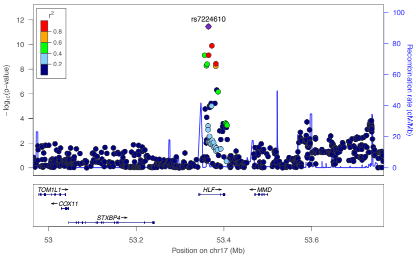


#### GTEx trans-eQTL DMD
#### brain nucleus accumbens basal ganglia

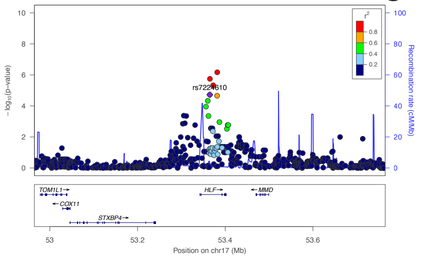


#### VEGFA locus
#### GWAS (Kottgen et al. 2013)

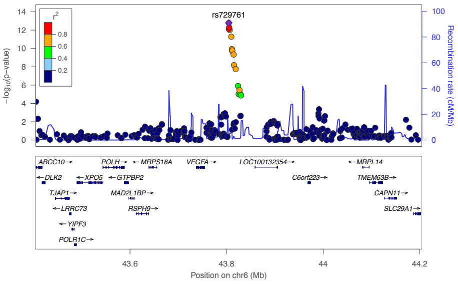


#### GTEx trans-eQTL CLPS
#### brain cerebellar hemisphere

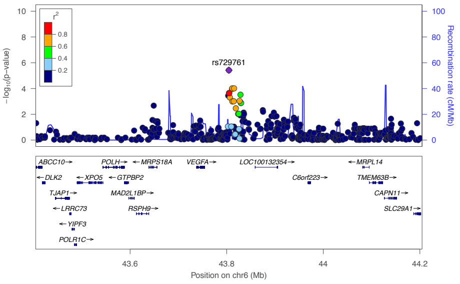

### Figure S7 cont.

#### MLXIP locus
#### GWAS (Kottgen et al. 2013)

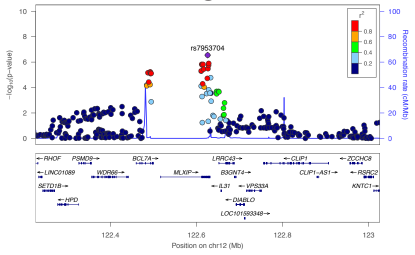


#### GTEx trans-eQTL NDUFA12
#### brain putamen basal ganglia

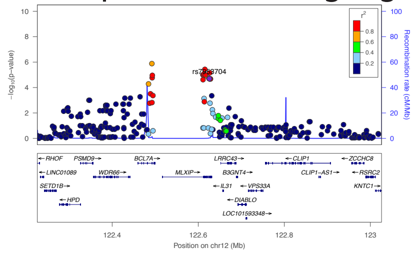


#### BCAS3 locus
#### GWAS (Kottgen et al. 2013)

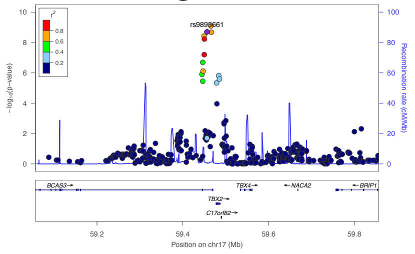


#### GTEx trans-eQTL TMEM117
#### prostate

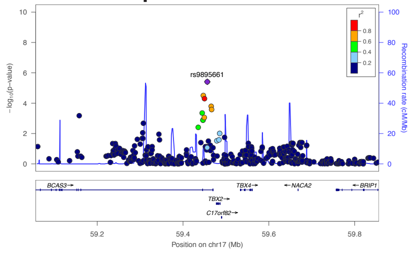


#### IDH2 locus
#### GWAS (Kottgen et al. 2013)

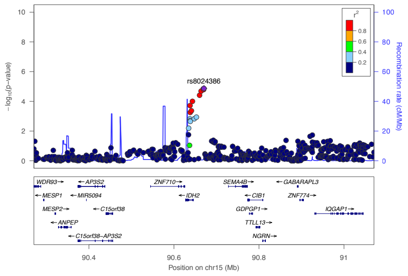


#### GTEx trans-eQTL MAPK6
#### brain amygdala

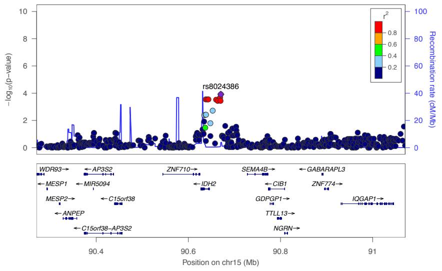


#### IDH2 locus
#### GWAS (Kottgen et al. 2013)

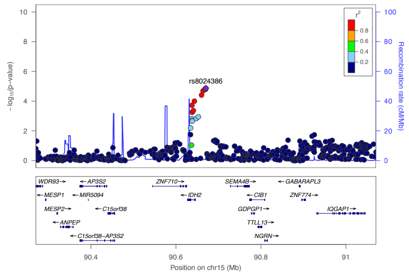


#### GTEx trans-eQTL ZBTB20
#### testis

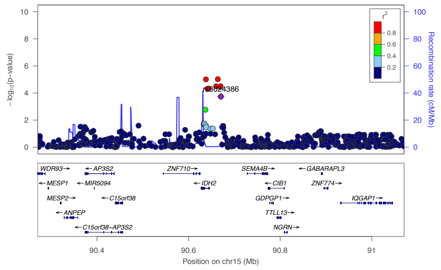

### Figure S8

#### UTRN locus
#### GWAS (Kottgen et al. 2013)

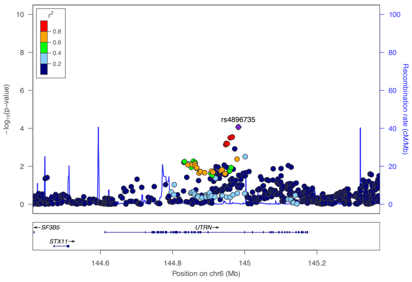


#### DMD locus
#### GWAS (Kanai et al. 2018)

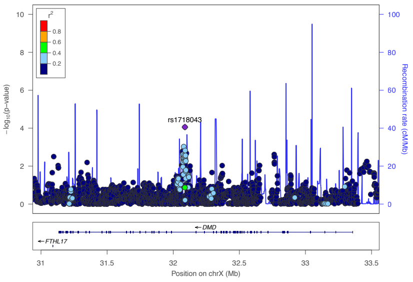

### Figure S9

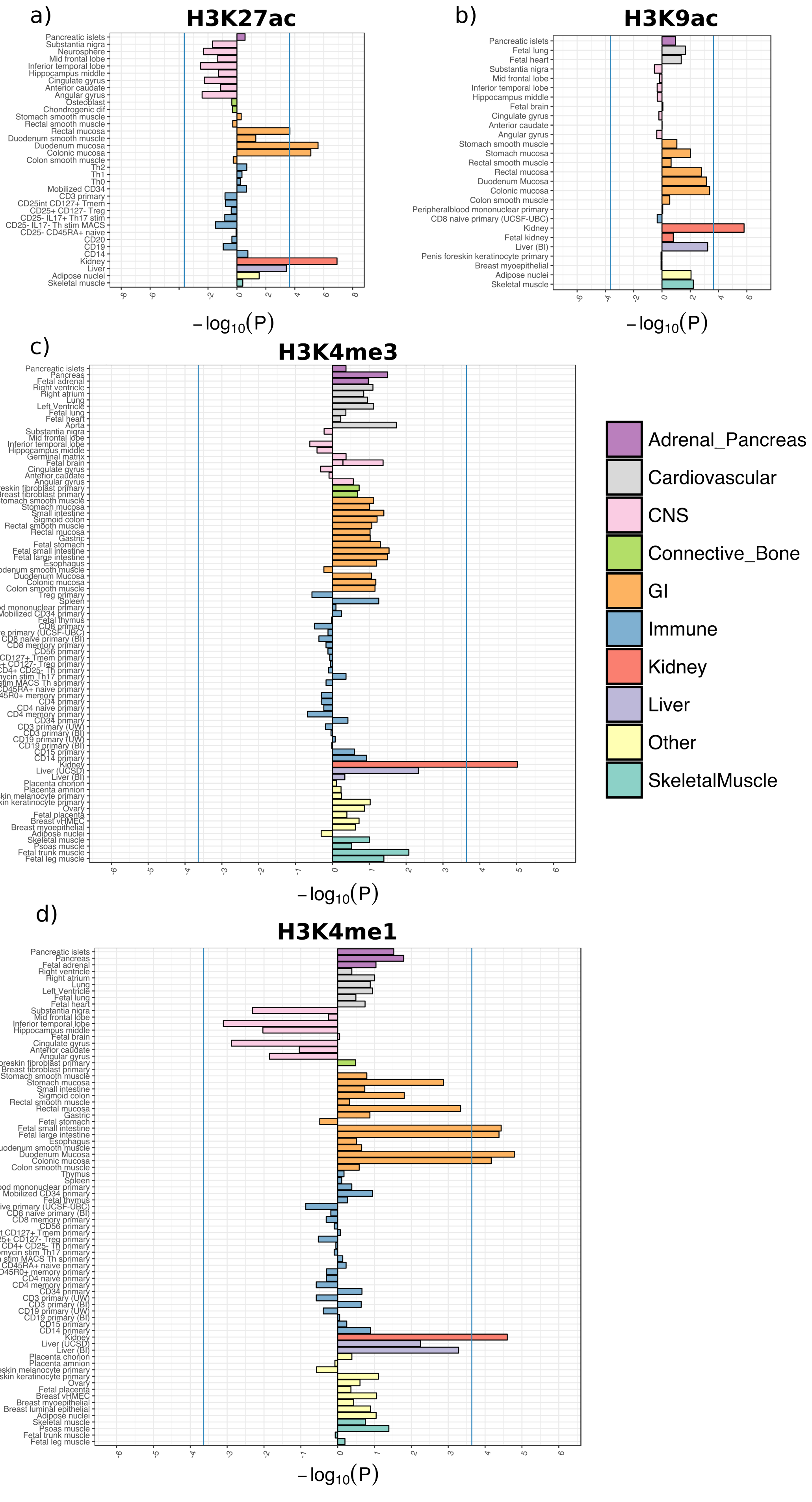
a)
H3K27ac
b)
H3K9ac
c)
H3K4me3
d)
H3K4me1
$-\log_{10}(P)$
Adrenal_Pancreas
Cardiovascular
CNS
Connective_Bone
GI
Immune
Kidney
Liver
Other
SkeletalMuscle

### Figure S10

#### SLC2A9 locus
#### GWAS (Kottgen et al. 2013)

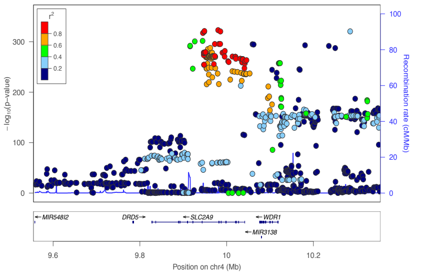


#### SLC2A9 locus
#### GWAS (Okada. 2012)

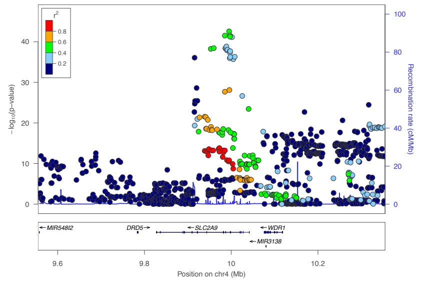


#### RP11-448G15.1 cis-eQTL
#### transformed lymphocytes

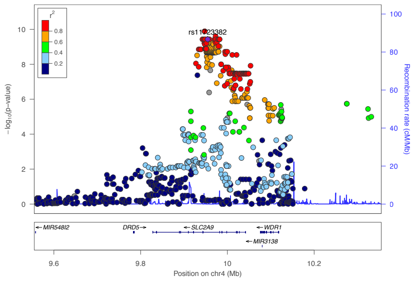


#### SLC22A12 locus
#### GWAS (Kottgen et al. 2013)

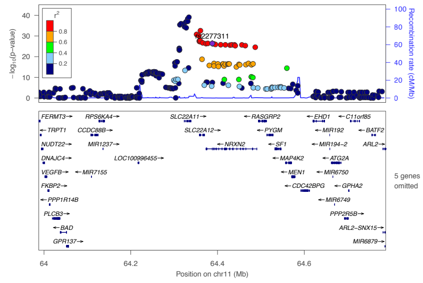


#### SLC22A12 locus
#### GWAS (Okada. 2012)

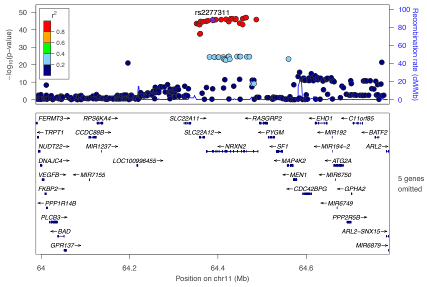


#### RNF169 trans-eQTL
#### heart atrial appendage

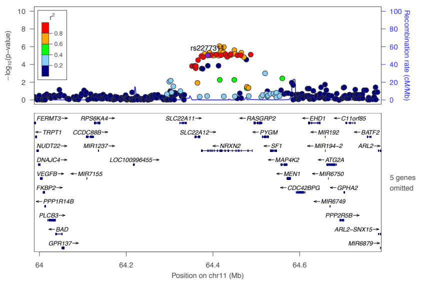
